# S^3^-CIMA: Supervised spatial single-cell image analysis for the identification of disease-associated cell type compositions in tissue

**DOI:** 10.1101/2023.03.17.533167

**Authors:** Sepideh Babaei, Jonathan Christ, Ahmad Makky, Mohammed Zidane, Kilian Wistuba-Hamprecht, Christian M. Schürch, Manfred Claassen

## Abstract

The spatial organization of various cell types within the tissue microenvironment is a key element for the formation of physiological and pathological processes, including cancer and autoimmune diseases. Here, we present S^3^-CIMA, a weakly supervised convolutional neural network model that enables the detection of disease-specific microenvironment compositions from high-dimensional proteomic imaging data. We demonstrate the utility of this approach by determining cancer outcome- and cellular signaling-specific spatial cell state compositions in highly multiplexed fluorescence microscopy data of the tumor microenvironment in colorectal cancer. Moreover, we use S^3^-CIMA to identify disease onset-specific changes of the pancreatic tissue microenvironment in type 1 diabetes using imaging mass cytometry data. We evaluated S^3^-CIMA as a powerful tool to discover novel disease-associated spatial cellular interactions from currently available and future spatial biology datasets.

## Introduction

The tissue microenvironment (TME) constitutes a spatially organized, dynamic and complex system of multiple cell types conferring emergent tissue properties in health and disease. Lymph nodes constitute a seminal example of how spatial organization of cell type composition in the TME confers emergent properties in terms of orchestrating the immune response to pathogens (1). In particular, dendritic cells and T cells are spatially matched in the lymph node’s T cell zone to increase the efficiency of adaptive immunity(2),(3). Until recently, it has been difficult to study such structures due to the technical requirement to both resolve spatial as well as high dimensional molecular profiles at the single-cell level. Moreover, the subsequent requirements to - typically computationally - integrate and interpret the resulting data to the end of defining descriptions or models of the TME and their disease-associated aberrations have been challenging.

This issue has been addressed technically with the development of high-dimensional proteomic, transcriptomic and epigenomic methods(4,5), single-cell spatial biology technologies, covering multiplexed fluorescence *in situ* hybridization (FISH), imaging mass cytometry (IMC) (6) and multiplexed microscopy such as cyclic immunofluorescence (CyCIF) (7) and co-detection by indexing (CODEX) (8,9). These approaches enable joint measurement of dozens of features in FISH, up to hundreds in multiplexed microscopy, and tens of thousands in spot-based spatial transcriptomics. Different resolutions, from local tissue spots as in spatial transcriptomics (e.g., high-definition spatial transcriptomics (10), Visium(11) and DBit-seq (12)) to subcellular resolution (e.g., FISH, IMC, CyCif, CODEX) are possible. Further, dedicated image processing, segmentation, and registration approaches have been developed to process and visualize the resulting data and quantify their respective signal intensities and distributions(13–15).

The resulting data lends itself for defining and mapping out TME arrangements. Two types of TME definitions can be distinguished: **unsupervised** definition of TME arrangements from the spatial single-cell data alone and **supervised** definition of TME characteristics associated with external cues such as disease state. While the former approaches allow for defining cell type composition of the TME, the latter approaches go a step further to identify differential cell type composition across conditions, including possibly novel, so far unappreciated cell subtypes. **So far, only unsupervised approaches of the first kind have been reported, and supervised approaches of the second kind are lacking**.

Approaches of the first kind include “Histology Topography Cytometry Analysis Toolbox” (histoCAT) (16) enabling unsupervised spatial interrogation of cell–cell interactions in IMC data. The neighborhood analysis in histoCAT examines if a certain cell type is located significantly closer to another cell type than expected by chance using two individual one-tailed permutation tests. “stLearn” (Spatial Transcriptomics Learn) was developed to calculate cell-cell interactions based on the unsupervised clustering of morphological similarity of spatial transcriptomics data (17). Briefly, stLearn utilizes unsupervised clustering to group similar spots into clusters (i.e., spatial morphological gene expression) and then significant cell-cell interactions are detected by a permutation approach. The tissue location hotspots are determined as locations in the tissue where there is both high interaction activity and diverse cell type co-localization. The “Giotto” analyzer and viewer (18) consists of tools to process and visualize spatial transcriptomics and proteomics expression data. The toolbox introduced new methods to identify a feature (gene or protein) that constructs a coherent spatial pattern based on unsupervised clustering and statistical enrichment of spatial network neighbors. In sum, these approaches often utilize nearest neighbors and other statistical approaches to identify relationships between different cell types and their TME based on an unsupervised strategy (19,20).

The above-mentioned methods are TME inference approaches of the first kind - as defined above - i.e., they are all unsupervised learning approaches that allow to define TME prototypes not allowing to infer TME differences associated and possibly explaining external cues such as disease state. Supervised learning approaches are required to overcome this limitation. Such approaches allow for identifying differences of TME models in comparative study settings, aiming at identifying differences of the same TME prototype across conditions, such as disease state or patient outcome. Supervised learning approaches enabling the detection of disease-associated TME compositions are so far lacking.

To close this gap, we present **supervised spatial single-cell image analysis (S**^**3**^**-CIMA)** for the identification of disease-associated cell type compositions in tissue microenvironments. S^3^-CIMA is a weakly supervised approach that leverages a single-layer convolutional neural network (CNN) architecture (21). We demonstrate the utility of S^3^-CIMA by identifying outcome-specific cell state compositions from a CODEX-based study of the colorectal cancer (CRC) TME. Specifically, we applied the method to a cohort of CRC patients to identify high-risk-specific spatial neighborhood cell type compositions in the tumor(20). Further, we use S^3^-CIMA to identify disease onset-specific changes of the pancreatic TME in type 1 diabetes in an IMC study (22). We expect that S^3^-CIMA will enable the study of other diseases and spatial biology data types and will generally be valuable to identify novel disease-associated cellular interactions.

## Results

### Weakly supervised learning of disease-associated TME compositions with S^3^-CIMA

We aim at learning (disease) condition-associated TME composition from multiparametric and spatially resolved single-cell data. Specifically, we consider the notion of condition-associated TME compositions as the spatial local enrichment of specific cell subsets with respect to a condition, i.e., *supervised spatial enrichment analysis* (**Fig. 1A**).

**Fig. 1.**
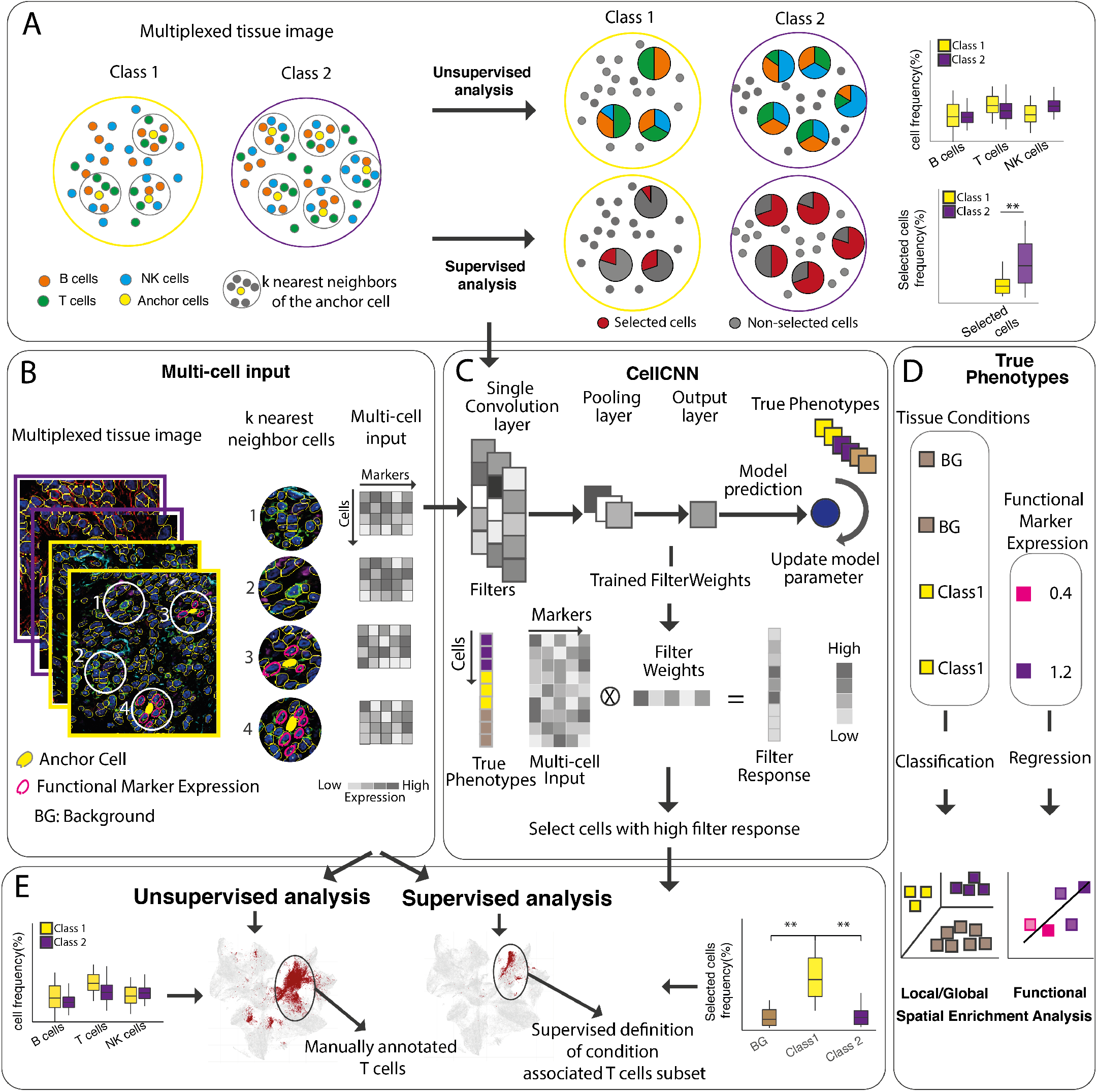
S^3^-CIMA overview and CRC global spatial enrichment analysis. S^3^-CIMA leverages a single-layer CNN architecture adopted from the CellCNN model (23). **(A)** Two types of TME definitions: unsupervised and supervised. Unsupervised approaches allow for defining cell type composition of the TME; unsupervised approaches compare the frequency of conventional cell types across the conditions, while supervised approaches identify differential subsets of cell type composition across tissue conditions. **(B)** The multi-cell input consists of k nearest neighbor cells around an anchor cell in a tissue image per phenotype label (i.e., class) and a set of marker expressions as the features. **(C)** The model is a single-layer CNN in which node activities are computed by weighted sums over marker expression values of each cell. The pooling layer summarizes the filter response of all cells in a multi-cell input per each convolutional filter. The trained filter weights are obtained by optimizing the CNN weights using true labels of multi-cell inputs. For each cell in the image data, the filter response is calculated by scalar product of the cell marker expression and the CNN trained filter weights. The cells with relatively high filter response indicate cells with a distinct signature to manifest the class of the image. The significant difference of frequency of selected cell populations between labels shows that the CNN model learns the difference between the phenotype labels. **(D)** Different types of supervised spatial enrichment analysis including local spatial enrichment analysis for identifying which cell subsets are enriched in the proximity of an anchor cell type, functional spatial enrichment analysis for identifying cell subsets enriched in the proximity of a specific functional activity of the tissue. **(E)** Two types of TME definitions: unsupervised and supervised. Unsupervised approaches allow for defining cell type composition of the TME, supervised approaches go a step further to identify differential cell type composition across tissue conditions, including condition-associated cell subtypes.

We distinguish three different types of *supervised spatial enrichment analysis*. In *global spatial enrichment analysis*, we aim at identifying which cell subsets are enriched in spatial neighborhoods across conditions, e.g., in CRC tumor tissues of disease manifestations with varying clinical outcomes. In *local spatial enrichment analysis* (anchor based) we instead aim at identifying condition-specific cell subset enrichment in the proximity of an *anchor cell type*, e.g., tumor-infiltrating cytotoxic T cells in CRC patients with different clinical outcomes. In *functional spatial enrichment analysis*, we aim at identifying cell subsets enriched in the proximity of a specific functional activity of the tissue, e.g., local expression of a functional immune cell marker.

The input of S^3^-CIMA is derived from high-dimensional *in situ* proteomic imaging data that has been processed up to single-cell segmentation, giving rise to a data matrix where every single cell (row) is associated with a profile of protein marker expression levels and spatial coordinates in the tissue image (columns) (**Fig. 1B**). The input for S^3^-CIMA are tuples of profiles of a set of spatially proximal cells, i.e., k-nearest *cell neighborhoods* (*k-NN*) around an anchor position (e.g., position of a cytotoxic T cell), associated with a phenotype indicating the originating condition (e.g., membership to a sample associated with a specific clinical outcome) (**Fig. 1B**). The association between these cell neighborhoods and their phenotype might be conferred by the occurrence of a cell subset not matching to a canonical cell type, or more generally, being unknown a priori. S^3^-CIMA addresses this challenge by a weakly supervised learning model, i.e., a CNN model(23) that takes these sets of cell profiles as input and learns their association with the phenotype label (**Fig. 1C**, see **Methods**).

Briefly, the model comprises a convolutional layer composed of filters, a pooling layer and a classification/regression output. The filters encode cell composition patterns from the molecular profiles (i.e., protein marker expression levels) by fitting the model to the training data. The pooling layer summarizes these pattern frequencies encoded by each filter, which in turn are used to associate with the classification/regression label (**Fig. 1D**).

The trained model is then used to identify the cell subsets critical for association with the phenotype label (e.g., patient outcome, treatment response, survival). Therefore, the trained filter weights correspond to molecular profiles and not to predefined annotated cell types of cell subsets in the local anchor cell neighborhood. These cell subsets are further characterized with respect to their cell type and molecular profile (**Fig. 1E**). The proximal enrichment of these cell subsets further motivates hypotheses about putative mechanisms conferring this enrichment (e.g., paracrine signaling).

We evaluated the S^3^-CIMA workflow for studies based on two different *in situ* proteomic imaging technologies, i.e., CODEX highly multiplexed fluorescence microscopy data of 56 markers in tumor tissues of 35 patients with low- and high-risk CRC, as well as IMC data of 35 markers in pancreas tissues from 12 donors with type 1 diabetes (T1D).

### S^3^-CIMA identifies spatially enriched cell subsets associated with differential CRC outcome

The spatial organization of the TME in CRC, indicating the colocalization and density of immune cells, is linked to disease progression and patient survival(24,25). Therefore, we applied S^3^-CIMA to the data of the CRC study(20), where patients were stratified in two groups with differential survival, i.e., 17 patients with Crohn’s-like reaction (CLR) and 18 patients with diffuse inflammatory infiltration (DII) (**Fig. 2A**). It has been previously reported that overall survival of patients with CLR is significantly longer than that of patients with DII(26). All cells in the CODEX images had been annotated to 28 unique cell types including 18 immune, 6 stromal, 2 mixed and 1 tumor cell groups(20) (**Fig. 2B, S1**). The study showed the differences in the frequency of immune cell compositions between CLR and DII patients in the individual TMEs. Most notably, CLR patients had higher frequencies of B cells and lower frequencies of macrophages than DII patients, respectively (**Fig. S2**). The study determined different cellular neighborhoods, i.e., tissue regions with a spatial localization of a cell type by means of an unsupervised framework, and showed that in the DII patients PD-1+ CD4+ T cells were enriched within a granulocyte cellular neighborhood. We expanded this analysis by assessing in an unbiased data-driven fashion the spatial enrichment of specific, possibly non-canonical cell subsets in the CRC TME between CLR and DII patients using the S^3^-CIMA model (**Fig. 2A**). We selected 100 random subsets of cells that are in k-NN (10 ≤ k ≤100) in each tissue image as the input for S^3^-CIMA. Each input was labeled according to the image group as CLR or DII (**Fig. S3**). The classification model was trained on 12 samples from each CRC group (24 staples in total) and tested on the remaining 11 samples (**Fig. 2C**). The trained model was used to identify the *selected cells*, i.e., the cell subset whose molecular profile (and not their annotated cell types, **Fig. 1A**) is relevant for classification of the CLR/DII patients (see **Methods**). The model robustly selected cells that are more frequent in the CLR group for neighborhood size k = 30 (p = 1.6e-12, Wilcoxon test) and in the DII group for neighborhood size k = 50 (p = 9.1e-13, Wilcoxon test) (**Fig. 2D, S5**).

**Fig. 2.**
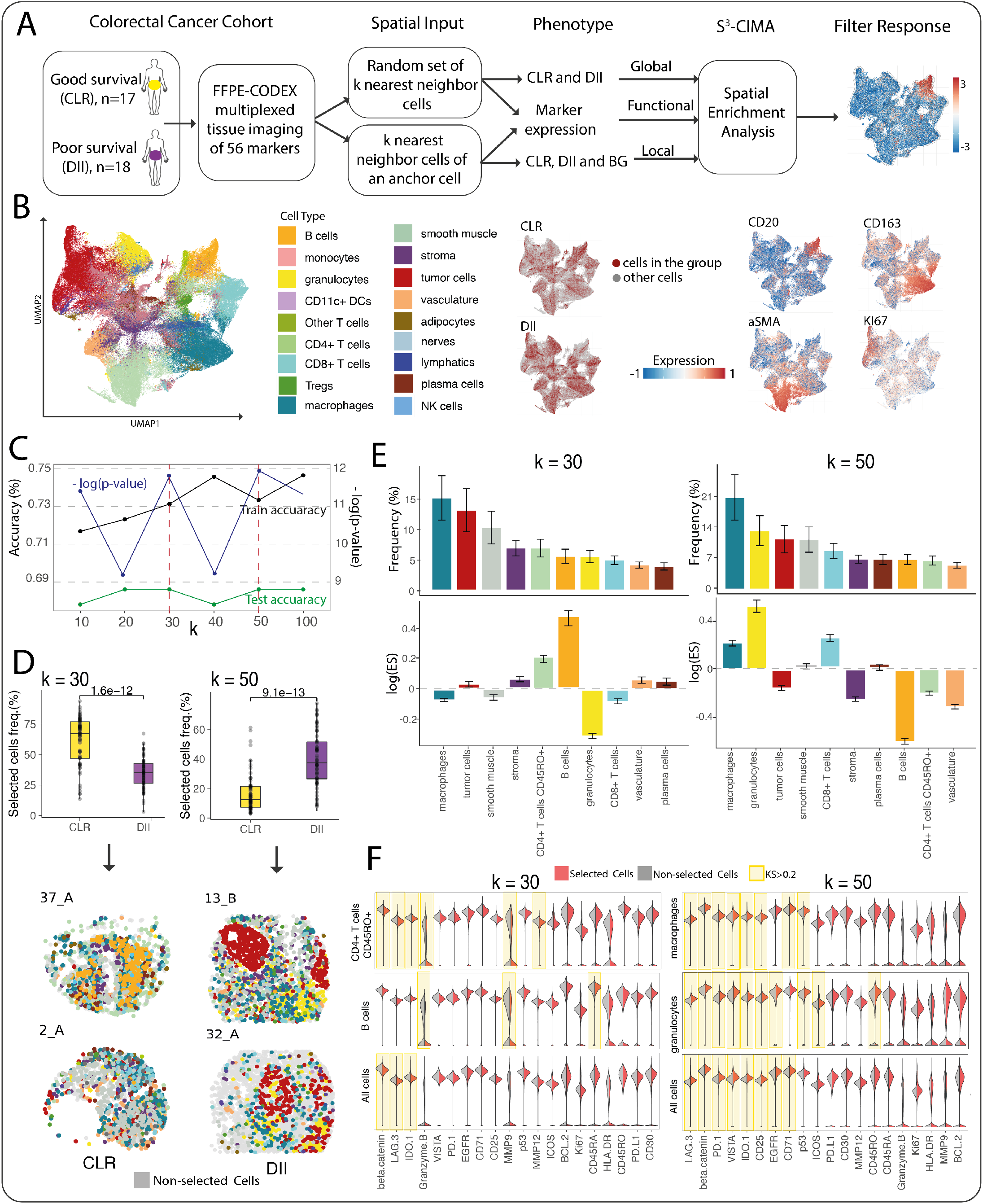
S^3^-CIMA global spatial enrichment analysis. **(A)** S^3^-CIMA global spatial enrichment analysis applied on the 140 CODEX images of 35 CRC patients. **(B)** Uniform Manifold Approximation and Projection (UMAP), indicating different cell types in the CRC tumor microenvironment. UMAP plots showing origin of single cells by clinical group of patients (CLR, DII) and expression of indicated markers. **(C)** S^3^-CIMA classification model performance (test and train accuracy) and -log(p-value) of comparing the frequency of selected cells by model across different cell neighborhood sizes (10 to 100). **(D)** The frequency of selected cells between groups in k = 30 (significantly more from CLR group) and k = 50 (significantly more from DII group) are shown. Selected cells (colored by cell type, color legend **Fig. 2B**) are mapped back to the corresponding patient CODEX images in both CLR and DII groups (also **Fig. S8-S11**). **(E)** The frequency and enrichment score (ES) of selected cells across the cell types. Bars are colored by cell type. A positive log value of ES indicates the enrichment of the corresponding cell type. **(F)** Density of the marker expression showing greatest differential abundance in terms of the Kolmogorov– Smirnov (KS) two-sample test statistics between the selected and non-selected cell subsets in all cell populations (regardless of the cell types) and between the selected and non-selected specific cell type subsets (also **Fig. S6, S7**). The high differential abundance (KS statistics) is highlighted by yellow color. The expression values were normalized between 0 and 1.

Since the selected cell subset consists of multiple cell types, we assessed the enrichment of each cell type of the selected cells by calculating an enrichment score (ES) across the patient groups, i.e., DII or CLR (see **Methods**). We found that at k = 30 the selected cells were predominantly from the CLR group and composed of macrophages (16%), tumor cells (14%), smooth muscle (10%), stroma (7%), and CD4+ T cells CD45RO+ (7%) (**Fig. 2E**). The selected CD4+ T cells CD45RO+ and B cells were significantly enriched (i.e., ES > 1) in the CLR group (**Fig. 2E**). For example, the selected B cells and CD4+ T cells CD45RO+ comprise a subset that exhibits significantly lower expression of Granzyme B compared to the non-selected cells (**Fig. 2F, S6**). We identified that the frequencies of subsets (i.e., selected cells by S^3^-CIMA) of B cells, CD8+ T cells, plasma cells, stroma and vasculature are significantly higher in the CLR than DII group (**Fig. S3**), while the aforementioned cell type frequencies are not significantly different when all cells are considered (i.e., unsupervised analysis) (**Fig. S2**). On the other hand, we found that at k = 50 the selected cells were predominantly from the DII group and composed of macrophages (21%), granulocytes (13%), tumor cells (12%), smooth muscle (12%) and CD8+ T cells (8%) (**Fig. 2E**). The selected immune cells have higher expression of the checkpoint molecules LAG3, PD-1 and VISTA compared to the bulk of their corresponding cell types that are in the tissue but not selected (**Fig. 2F, S7**). The S^3^-CIMA model can thus identify specific cell subsets at different neighborhood sizes which associate with either poor or superior surviving patients (i.e., at k=30 and k=50). This suggests that S^3^-CIMA can capture the disease-associated TME characteristics at multiscale cellular neighborhoods. We also identified that the frequencies of subsets of tumor cells, CD4+ T cells CD45RO+, stroma and CD68+ macrophages which are selected by S^3^-CIMA compared with non-selected cells from the same cell types are significantly higher in the DII than in the CLR group (**Fig. S4**). However, the frequencies of these cell types are not significantly different when all cells are considered (i.e., unsupervised analysis) (**Fig. S2**).

### S^3^-CIMA reveals high enrichment of PD-1+ CD4+ T cell subset in granulocyte neighborhoods of DII patients

Schürch et al.(20) defined nine cellular neighborhood classes as tissue regions with a high density of a specific cell type and assessed if the frequencies of these exhibit any differences between two CRC patient groups. This analysis showed that except for the follicle cellular neighborhood which was highly enriched in CLR patients, none of the other cellular neighborhood categories differed significantly. It also identified that PD-1+ CD4+ T cells enrichment in the granulocyte neighborhood correlated with overall survival in the DII patient group. The study therefore suggested that to understand the underlying process shaping the CRC TME, cell types and the cellular neighborhood should be considered simultaneously.

S^3^-CIMA allows to achieve this goal in an unbiased data-driven fashion, by learning disease associated cellular neighborhoods around specific anchor cell types, i.e., the *anchor cell niche* (**Fig. 1A**). We considered each cell type in the dataset as the anchor cell type. We generated condition-specific (i.e., CLR or DII) CNN models with the adequate classification accuracy when using B cells, CD4+ T cells, tumor cells, M2 macrophages and granulocytes as anchor cell types across a range of k-NN sizes (10 ≤ k ≤50) (**Figs. S12**).

Considering granulocytes as the anchor cell type, we found that the frequency of selected cells is significantly higher in the DII group compared to CLR and background at k = 30 (p = 4.6e-5 and p = 1.2e-15, Wilcoxon test, respectively) (**Fig. 3A, S13**) and composed of granulocytes (58%), macrophages (7%), tumor cells (6%) and smooth muscle (5%) (**Fig. 3B**).

**Fig. 3.**
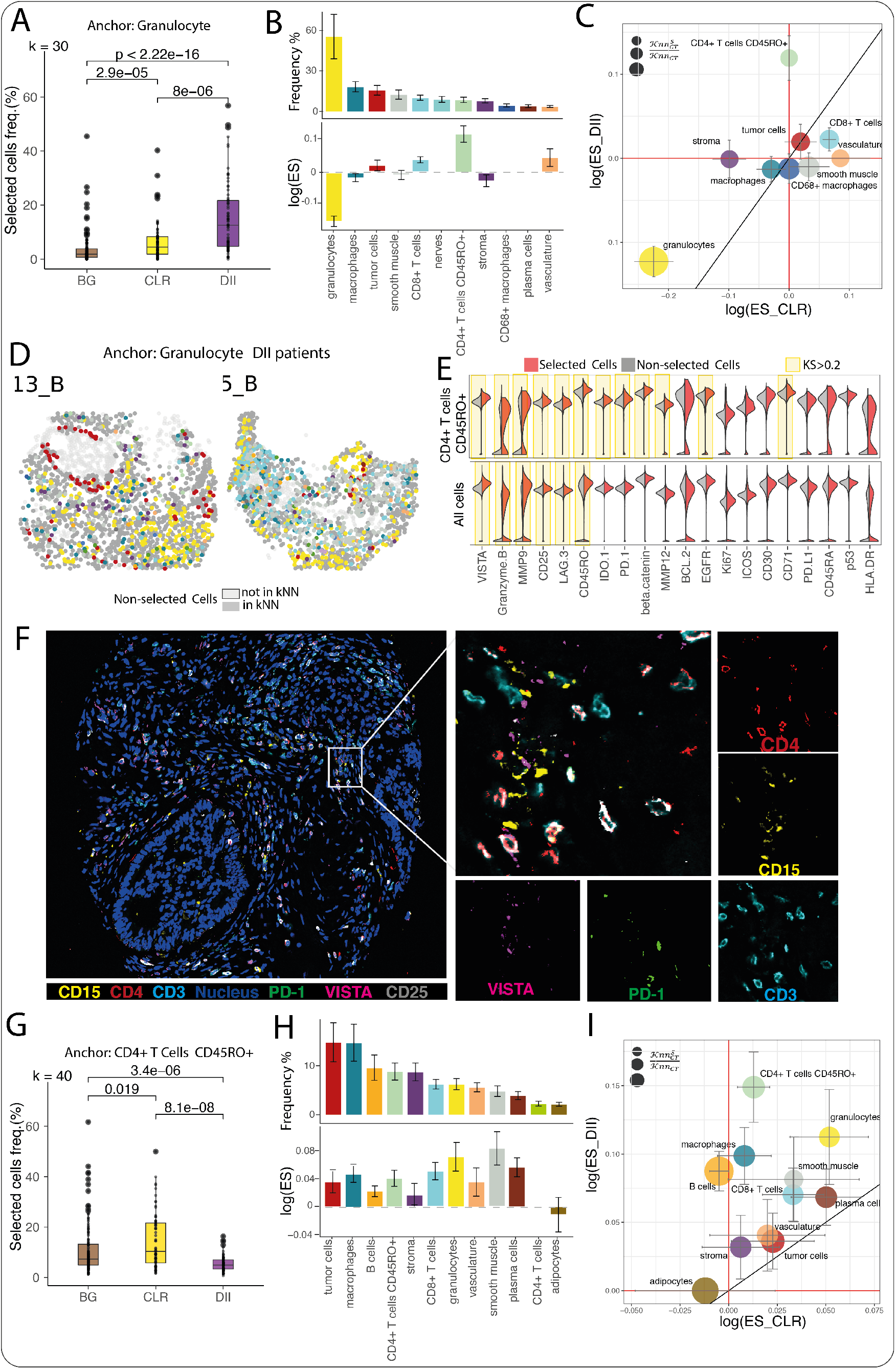
S^3^-CIMA local (anchor based) spatial enrichment analysis. The multi-cell inputs generated from k nearest neighbor cells around granulocytes (k = 30) and CD4+ T cells CD45RO+ (k = 40) across all 35 CODEX images of the CRC dataset. Each multi-cell input was labeled by a CRC outcome class (CLR or DII) and background (BG) for the classification task. **(A)** Frequency of selected cells across the patients between groups with granulocytes as an anchor. The cells with the high filter response have a significantly increased frequency in the DII group **(B)** Frequency and ES of selected cells across the cell types obtained from local enrichment spatial analysis for granulocytes as the anchor. Bars are colored by cell type. A positive log value of the ES indicates the enrichment of the corresponding cell type. **(C)** Bubble plot of ES of each cell type in the proximity of granulocytes comparing different survival groups (median of ESs across all patients). The MAD is shown by the error bar. The color and size of the bubbles indicate the cell types (color legend **Fig. 2b**) and the ratio of the number of selected cells to the total number of cells of the specific type, respectively. **(D)** Selected cells (colored by cell type, color legend **Fig. 2b**) are mapped back to the corresponding patient CODEX images in the DII groups (**Fig. S14-15**). **(E)** Density of functional marker expression showing greatest differential abundance in terms of the KS two-sample test statistics between the selected and non-selected cell subsets in all cell populations (regardless of the cell types) and between the selected and non-selected specific cell type subsets (also **Fig. S16**). The expression values were normalized between 0 and 1. **(F)** Codex image of Patient 30 (DII group). The spatial neighborhood of one granulocyte as the anchor is zoomed in. The PD-1, CD4, CD15 and CD68 markers intensities of the selected cells around this anchor are shown. **(G)** Frequency of selected cells across the patients between groups with CD4+ T cells CD45RO+ as an anchor. The cells with the high filter response have significantly increased frequency in the CLR group. **(H)** Frequency and ES of selected cells across the cell types obtained from local enrichment spatial analysis for CD4+ T cells CD45RO+ as the anchor. Bars are colored by cell type. A positive log value of the ES indicates the enrichment of the corresponding cell type. **(I)** Bubble plot of ES of each cell type in the proximity of CD4+ T cells CD45RO+ comparing different survival groups (median of ESs across all patients). The MAD is shown by the error bar. The color and size of the bubbles indicate the cell types (color legend **Fig. 2B**) and the ratio of the number of selected cells to the total number of cells of the specific type, respectively.

To assess whether the selected cell types are spatially enriched in the granulocyte neighborhood and are not selected because of the abundance, we calculated an analytical ES for each cell type. The total number of cells of a given annotated cell type in the granulocyte neighborhood does not correlate with a higher ES (i.e., ES >1). The granulocyte niche is enriched for CD4+ T cells CD45RO+ (ES = 1.14), vasculature (ES = 1.05) and CD8+ T cells (ES = 1.04) (**Fig. 3B**). Interestingly, we confirmed that the granulocyte neighborhood is highly enriched for CD4+ T cells CD45RO+ in the DII group compared to the CLR group (**Fig. 3C**). To characterize the selected cell subset compared to non-selected cells and assess its differential functional capacity, we performed differential marker expression analysis. Specifically, we compared the functional marker expression, such as apoptosis, activation/proliferation, inhibition and cytokine signaling markers, in the selected subsets of cells versus non-selected cells within each specific cell type compartment. The selected cells in the granulocytes niche highly express VISTA (KS = 0.48), Granzyme B (KS = 0.32), and MMP9 (KS = 0.31) (**Fig. 3E, S16**). We also identified that the spatially enriched CD4+ T cells CD45RO+ subset in the granulocyte niche over-expresses T cell exhaustion/activation markers VISTA (KS = 0.44), PD-1 (KS = 0.22) and EGFR (KS = 0.28) (**Fig. 3E**). To confirm the protein marker activity of enriched cell subset in the granulocytes niche, we then mapped the selected cells in the granulocyte niche to the CODEX images and observed a high expression of PD-1 and CD4 markers in selected cell subsets (**Fig. 3D, F, S14-15**).

Then, to investigate the mutual interaction of CD4+ T cells CD45RO+ and granulocytes, we performed S^3^-CIMA local enrichment analysis considering CD4+ T cells CD45RO+ as the anchor cell type. We found that the frequency of selected cells is significantly higher in the CLR group compared to DII and background at k = 40 (p = 8.1e-8 and p = 0.019, Wilcoxon test, respectively) (**Fig. 3G, S13**) and composed of mostly tumor cells (15%), CD163+ macrophages (15%) and B cells (8%) (**Fig. 3H, S17-19**). The enrichment analysis showed that the CD4+ T cells CD45RO+ niche is enriched by smooth muscle (ES = 1.08), granulocytes (ES = 1.07), plasma cells (ES = 1.06) and CD8+ T cells (ES = 1.05) (**Fig. 3I**).

We then calculated the ES for each CRC group separately and interestingly, we also identified the spatially enriched granulocyte subset in the CD4+ T cells CD45RO+ niche which is enriched in the DII group, indicating a specific mutual interaction of these two cell types (**Fig. 3I**). This interaction can be confirmed by colocalization of representative cells in the original CODEX images (**Fig. 3F**). These results suggest that S^3^-CIMA can determine the local enrichment of the subset of a conventional cell type in the spatial neighborhood of a specific cell type as well as the one-to-one cell type interaction.

### Functional spatial enrichment analysis reveals an EGFR-expressing cell subset in the proximity of granulocytes

We then performed *functional spatial enrichment analysis* by S^3^-CIMA to identify cell subsets enriched in the proximity of an anchor cell, i.e., granulocytes (the same anchor that we used for the local enrichment analysis) varying in local epidermal growth factor receptor (EGFR) expression. EGFR plays a key role in different cellular functions, such as proliferation, apoptosis and differentiation. It has been shown that high expression of EGFR is common in many tumors. In particular in CRC, high expression of EGFR is associated with a poor prognosis (27).

To determine the effect of localized EGFR expression, we considered the phenotype label of a multi-cell input (i.e., cells in the k nearest neighborhood) as the average expression of EGFR over all k cells in the nearest neighborhood of each granulocyte, and then trained a regression S^3^-CIMA model (R2-score = 63.8%, RMSE = 0.28). To summarize the distribution of selected cells with respect to local EGFR expression, we report selected cell frequencies for either high/low, i.e., higher/lower than average EGFR expression across all considered cell neighborhoods. The model learns the spatial characteristics of both groups with two discriminative filters. We found that the frequency of selected cells is significantly higher in the EGFR^Low^ group compared to EGFR^High^ for filter 1 and vice versa for filter 2 at k = 20 (p = 2.2e-16, Wilcoxon test) (**Fig. 4A**). Considering the response of filter 1, we identified that the granulocyte niche is highly enriched in adipocytes, plasma cells and CD8+ T cells in the EGFR^Low^ group (**Fig. 4B, C**). The selected cell subset in the granulocyte niche is associated with significantly reduced EGFR, CD25 and Ki67 expression (**Fig. 4D**). The response of filter 2 determined that the granulocyte niche is highly enriched by B cells and tumor cells in the EGFR^High^ compared to the EGFR^Low^ group (**Fig. 4B**). The spatially enriched B cells subset in the granulocyte niche significantly over-expresses the interleukin-2 receptor CD25 (KS = 0.35), EGFR (KS = 0.34) and PD-L1 (KS = 0.32) (**Fig. 4D**). This result shows that colocalization of granulocytes and B cells is associated with high expression of EGFR in the TME. With this case, we demonstrated that S^3^-CIMA is capable of identifying determinants of niche composition that are associated with local signaling activity in the tissue such as EGFR signaling (**Fig. 4E**).

**Fig. 4.**
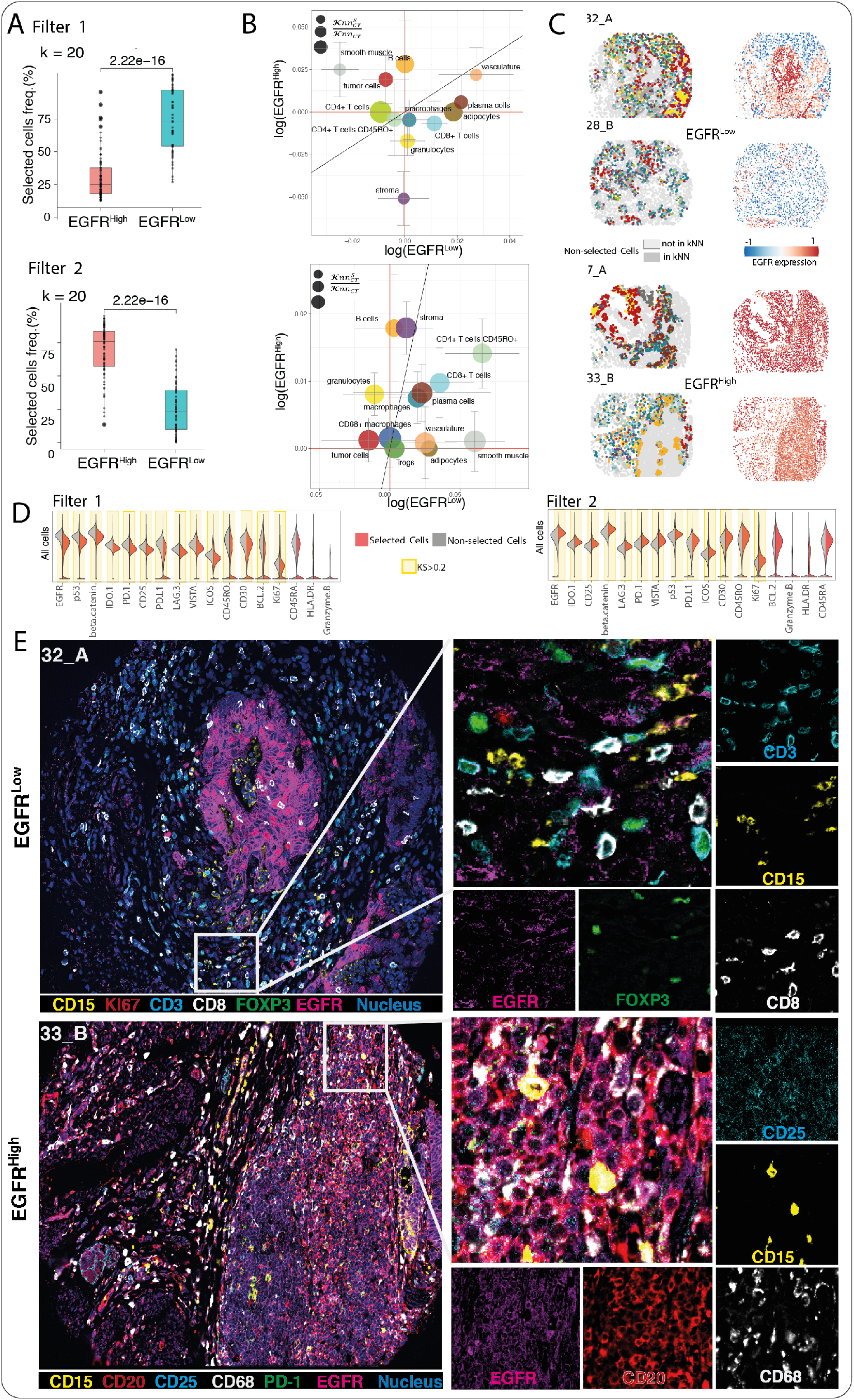
S^3^-CIMA functional spatial enrichment analysis. **(A)** The multi-cell inputs generated from k nearest neighbor cells around granulocytes (k = 20) across all 35 CODEX images of the CRC dataset. Each multi-cell input was labeled by the average of EGFR marker expression of the cells in that multi-cell input (continuous value) as a surrogate for local signal transduction activity for the regression task. Frequency of selected cells of functional enrichment analysis with granulocytes as an anchor comparing two groups of patients with high and low EGFR marker expression. The cells with a high filter response have a significantly increased frequency in the EGFR^Low^ group in filter 1 and EGFR^High^ group in filter 2. **(B)** Bubble plot of enrichment score of each cell type in the proximity of granulocytes showing the cytokine signaling activity of the EGFR marker in each group. The MAD is shown by the error bar. The color and size of the bubbles indicate the cell types (color legend Fig.2b) and the ratio of the number of selected cells to the total number of cells of the specific type, respectively. **(C)** Selected cells (colored by cell type, color legend Fig.2b) and the EGFR expression value for each cell are mapped back to the corresponding patient CODEX images in both EGFR^High^ and EGFR^Low^ groups (also **Fig. S20-S23**). **(D)** Density of functional marker expression showing greatest differential abundance in terms of the Kolmogorov–Smirnov two-sample test between the selected and non-selected cell subsets in all cell populations and per cell types for regression task. The expression values were normalized between 0 and 1. **(E)** Codex images of Patient 16 (Spot 32_A, DII group, EGFR^Low^) and Patient 17 (Spot 33_B, CLR group, EGFR^High^). The spatial neighborhood of one granulocyte as the anchor is zoomed in. The mentioned marker intensities of the selected cells around this anchor are shown.

### S^3^-CIMA reveals high enrichment of beta cells in cytotoxic T cell neighborhoods in type 1 diabetes onset patients

We examined if S^3^-CIMA is also capable of finding a spatial pattern of cells in IMC data. To accomplish this, we used a type 1 diabetes (T1D) study including IMC measurement of 35 protein markers of pancreas tissues from 12 donors, comprising healthy-, T1D disease onset, and long disease duration conditions (22) (**Fig. 5A**). T1D is an autoimmune disease which is caused by an immune system attack on insulin-producing beta cells in the pancreatic islets of Langerhans. The spatial interaction between islets and immune cells can be involved in the progression of the disease (28). This study demonstrated that beta cell destruction is preceded by recruitment of cytotoxic and helper T cells in T1D disease onset.

**Fig. 5.**
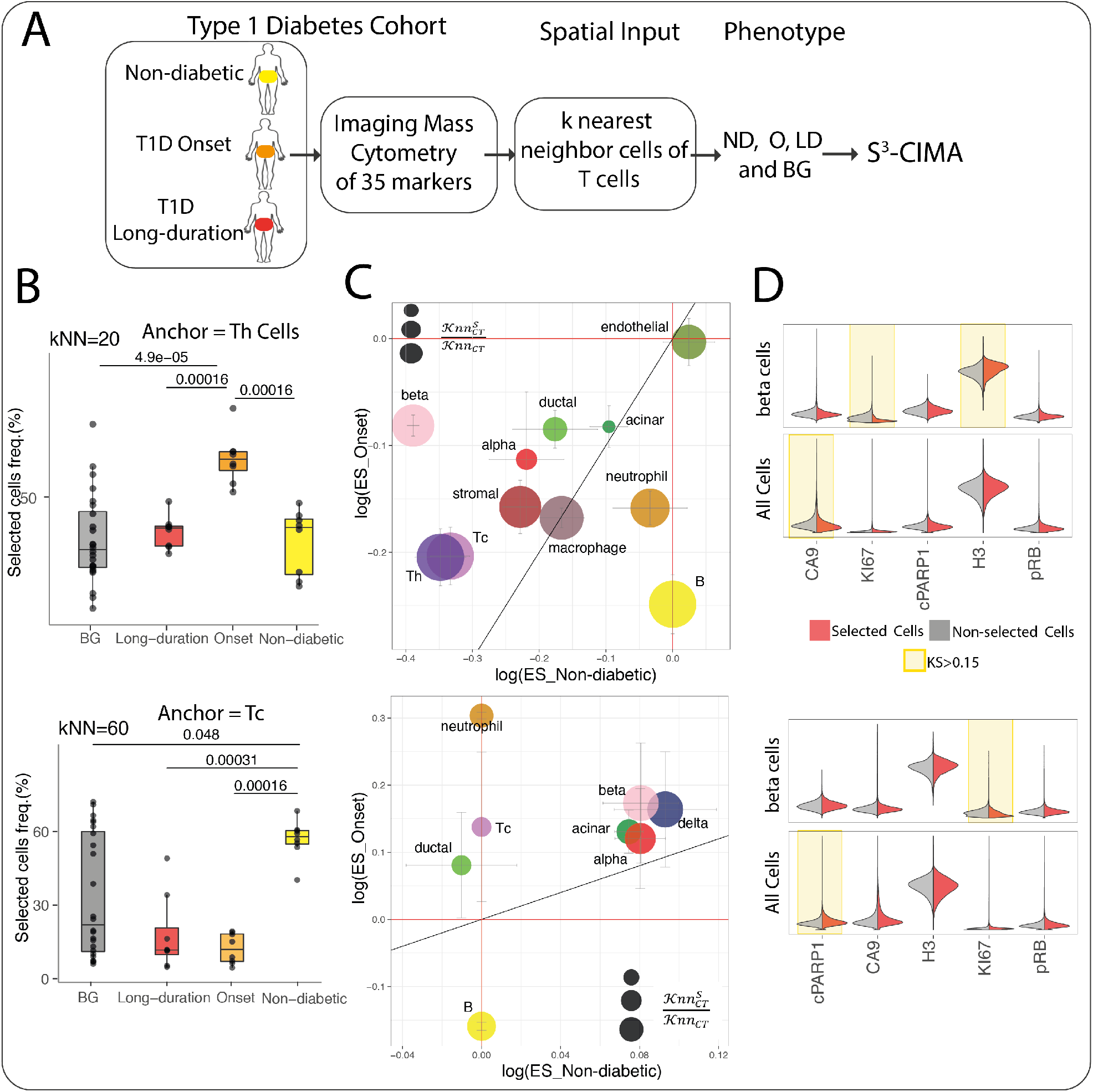
S^3^-CIMA local (anchor based) spatial enrichment analysis. **(A)** S^3^-CIMA local spatial enrichment analysis was applied on IMC measurement of 35 protein markers of pancreas tissues from 12 human donors, comprising healthy-, T1D onset- and long disease duration conditions. **(B)** Frequency of selected cells between disease stages with helper T cells as an anchor. The cells with the high filter response have significantly increased frequency in the onset T1D group. Frequency of selected cells between disease stages with cytotoxic T cells as an anchor. The cells with the high filter response have significantly increased frequency in the non-diabetic group. **(C)** Bubble plot of ES of each cell type in the proximity of helper T cells shows that there is no high enrichment of the presence of any cell type. The color and size of the bubbles indicate the cell types and the ratio of the number of selected cells to the total number of cells of the specific type, respectively. Bubble plot of ES of each cell type in the proximity of cytotoxic T cells shows that there is a high enrichment of the presence of beta cells from the onset T1D group. **(D)** Density of functional marker expression showing greatest differential abundance in terms of the Kolmogorov–Smirnov two-sample test between the selected and non-selected cell subsets in all cell populations and beta cells. The expression values were normalized between 0 and 1.

We performed S^3^-CIMA analysis for local spatial enrichment analysis to understand association between islet cells and immune cells during T1D progression. The study showed that the association between immune cells and islet cells is not common; however, during T1D onset, because of the assembly of a destructive immune reaction, cytotoxic and helper T cells are recruited to beta cell-rich islets (29). Similarly, we also observed that the presence of immune cells (monocytes, neutrophils and B cells) in spatial proximity of islet cells (alpha, beta, gamma and delta cells) is not frequent in all three disease stages, i.e., there is no spatially enriched subset of islet cells when any of these immune cell types is an anchor cell. However, we observed that when we performed S^3^-CIMA with T cells as an anchor (T helper cells, cytotoxic T cells (**Fig. 5B**) and naïve T cells (**Fig. S24**)), there is a high enrichment of the presence of beta cells in the neighborhood of cytotoxic T cells from the T1D onset group (**Fig. 5C, D**). It is known that cytotoxic and helper T cells are involved in the destruction of beta cells in T1D (30).This result confirms the previous finding of (22) that showed that beta cell destruction is preceded by recruitment of cytotoxic and helper T cells in T1D disease onset. In summary, we find that S^3^-CIMA is capable of discovering novel disease-associated spatial cellular interactions from various spatial biology data types.

## Discussion

Understanding how cell type compositions in the TME differ from one disease condition to another can pinpoint tentative disease mechanisms. S^3^-CIMA provides a *supervised spatial enrichment analysis* by learning from disease-associated TME composition to systematically identify spatially enriched cell subsets associated with the disease conditions. We showed that S^3^-CIMA achieves this goal in an unbiased data-driven fashion, by learning disease associated cellular neighborhoods around specific anchor cell types (*local spatial enrichment analysis*) or random sets (*global spatial enrichment analysis*). We demonstrated, for both CODEX and IMC data, that S^3^-CIMA can reveal novel subpopulation structures and tissue organization possibly missed by single modalities or other methods. The frequencies of selected subpopulations are significantly different across the phenotypes (e.g., CLR and DII groups in the CRC cohort analyses), while the frequency of the whole population of those cell types (i.e., unsupervised analysis) are similar in the phenotype groups (**Figs. S2-4**). Identified cellular subpopulations can, in principle, always be further subdivided to reflect a finer characterization of observed heterogeneity. Subdivision based on multiple modalities provides an opportunity to identify more meaningful biological distinctions.

Here, we examined the S^3^-CIMA model on antibody-based proteomic imaging data that is inherently limited to sub-genome-wide analyses. This limited spectrum of molecular parameters possibly precludes the detection of sample condition-associated cell subsets defined by unmeasured molecular markers. However, the increase in spatial resolution of spatial transcriptomic approaches is likely going to alleviate this limitation and allow S^3^-CIMA to comprehensively assess the cell states across genome-wide expression profiles. Further, the greedy nature of the model fitting can fail to report the cell subsets that are less but still significantly correlated with the supervision signal than the selected cell subset.

Up to now, most of the spatial proteomic imaging data analysis efforts applied unsupervised methods that cannot use the associating external cues of interest directly. However, an increasing number of spatial proteomic imaging studies in the field of health and disease are and will be comparative, i.e., aiming at identifying spatial cell type composition patterns that explain external cues, such as physiological state, disease state, therapy response or signal transduction activity. We showed that S^3^-CIMA can use both discrete and continuous states of these external cues as the sample phenotypes. S^3^-CIMA constitutes a machine learning model that addresses this issue and - as first of its kind - is expected to enable productive interpretation of such comparative studies in the future.

## Materials and Methods

### Notation used throughout the paper

|  |  |
| --- | --- |
| C | Multi-cell input |
| (X, Y) | Position of a cell |
| k | Number of nearest neighbor cells |
| T | Cell type |
| p | p-value |
| k-NN | k-nearest neighbors |
| KS | Kolmogorov–Smirnov test statistic |

### Acronyms used throughout the paper

|  |  |
| --- | --- |
| CRC | colorectal cancer |
| CLR | Crohn's-like reaction |
| DII | diffuse inflammatory infiltration |
| TME | tissue microenvironment |
| T1D | type 1 diabetes |
| ES | enrichment score |
| CNN | convolutional neural network |
| BG | background |
| MAD | median absolute deviation |
| EGFR | epidermal growth factor receptor |

### Datasets

#### Colorectal cancer

Here, we used the data of an already published CRC cohort including 56-marker multiplexed CODEX tissue imaging data of 35 CRC patients including 140 TMA spots i.e., 4 CODEX images for each patient (20). The marker set includes lymphocyte and myeloid cell canonical markers, functional immune cell markers (e.g., cytokine signaling, activation / proliferation, inhibition / checkpoint markers) and several auxiliary markers.

The flat table of the images was used for computing the spatial input of S^3^-CIMA indicating 258,385 cells across 140 CODEX images of 35 patients as rows and patient ID, marker intensity, position (i.e., X/Y coordinates), survival group (i.e., DII or CLR) and annotated cell type (29 cell types) as columns. Marker intensity values were log transformed by adding an offset (log(1e-3+ x)) and scaled by z-score across the dataset prior to calculating spatial inputs (We followed the same preprocessing procedure of (20)). The spatial inputs of 24 patients (12 patients from each survival group) were randomly split into training and validation sets (80% and 20% respectively). The spatial input of the remaining 11 patients were used as the test. We calculated the nearest neighbors and enrichment analysis per TMA spots (i.e., images).

#### Type 1 diabetes

The type 1 diabetes (T1D) cohort includes IMC measurement of 35 protein markers of pancreatic islets of Langerhans tissues from 12 human donors, comprising healthy (n=4), recent-onset T1D (<0.5 years, n = 4) and long-standing T1D duration (>8 years, n = 4)(22). The data includes two sections originating from different anatomical regions of the pancreas (tail, body, or head) for each donor. The flat table of the IMC images was used for computing the spatial input of S^3^-CIMA indicating 1,776,974 cells across 845 IMC images of 12 patients as rows and patient id, slide id (24 slides), cell position in each image (i.e., X/Y coordinates) cell category (exocrine, immune, islet, other, unknown), cell type (16 cell type) as columns. Marker intensity values were arcsinh transformed and scaled by z-score across the dataset prior to calculate spatial inputs. The spatial inputs of 9 patients (3 patients from each disease condition group) were randomly split into training and validation sets (80% and 20% respectively). The spatial input of the remaining 3 patients were used as the test set.

#### S^3^-CIMA model training

S^3^-CIMA implements a weakly supervised CNN model to identify cell subsets whose frequency distinguishes the considered phenotype labels (i.e., disease associated conditions). The model is adopted from the CellCNN model (23), comprising a single layer CNN, a pooling layer and a classification or regression output, and using groups of cell expression profiles (multi-cell inputs) as input. S^3^-CIMA uses multi-cell inputs, which are generated k nearest cell neighbors in a tissue image per class label with a set of marker expressions as the features (*global spatial enrichment analysis)*. For *local spatial enrichment analysis*, S^3^-CIMA uses multi-cell inputs as *k* nearest cell neighbors around anchor cells of a specific cell type. Training and testing of the model are performed as described in (23). Briefly, 200 models were trained and the model with highest predictive accuracy on the validation set was selected.

#### Global spatial enrichment analysis

Global spatial enrichment analysis aims at identifying cell subsets that are spatially co-localized across the multiple conditions. The multi-cell input is generated using nearest neighbor cells of *N* randomly selected cells in each tissue image with a set of marker expressions as the features. A multi-cell input is the set of marker expression vectors of the k nearest neighbor cells [**C**_*i*_]_*KM*_, i ∈ {1, 2,…,*N*}, where *K* and *M* are the number of cells and number of markers, respectively.

We performed S^3^-CIMA global spatial enrichment analysis of the CRC cohort by selecting *N* random cells in each patient CODEX image and calculating k-NN (10 ≤ k ≤100) based on the Euclidean distance. Each **C**_*i*_ input was labeled according to its corresponding image group as CLR or DII. Since some TMA images include only 200 cells, we selected *N* = 100 random cell subsets as the spatial multi-cell inputs of the classification model.

#### Local spatial enrichment analysis

Local (anchor based) spatial enrichment aims at identifying cell subsets that are enriched in spatial proximity of a specific anchor cell type across the multiple conditions. The multi-cell inputs k nearest neighbors of an anchor cell are denoted [**C**_*i*_]_*KM*_, i ∈ {1, 2,…*N*}, where *N, K* and *M* are the number of the anchor cell, number of neighboring cells and number of markers, respectively. We added a set of randomly selected k nearest neighbor cells as the background (BG) label to ensure that the cell signatures identified by the S^3^-CIMA indicate the anchor cell specific spatial colocalization.

#### Functional spatial enrichment analysis

Functional spatial enrichment analysis aims at identifying cell subsets enriched in the proximity of a specific functional activity of a specific anchor cell type, e.g., local proliferative activity of tumor cells. We examined the S^3^-CIMA ability to detect spatial enriched functional subset using the colorectal cancer dataset. The preprocessing and computing of the multi-cell input [C_*i*_]_*KM*_ is similar to the *local spatial enrichment* analysis. However, the multi-cell inputs were labeled according to the specific functional activity i.e., the average expression of a specific marker over all cells in [C_*i*_]_*KM*_. Here, the k-NN of that anchor cell type were computed, and the M-1 marker intensities of those selected k-cells were the features. The average of the functional marker intensities of the multi-cell inputs were used to capture the functional activity as the dependent variable during training of the model.

#### Quality of prediction of the regression model

To measure the accuracy and the predictive ability of the training, we used the *R*^2^ score and RMSE, respectively. The *R*^2^ score, shows the amount of variance of the true values, which can be explained by the independent variable, and therefore gives a probability about how well unseen samples can be predicted. It is defined as:

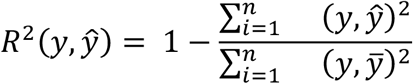

where *y* is the true value and *Ŷ* is the predicted value of the sample from a total of *n* samples. The term 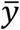 is the average of the true values.

The measure of RMSE, gives an idea of the standard deviation of the predictive error, which is the difference between the observed and predicted values. RMSE captures how spread out these values are. RMSE is obtained by taking the square-root of the mean squared error (MSE), another common metric for measuring the accuracy of the model.

#### Characterization of spatially enriched cell subsets

The downstream analysis using the trained filter weights facilitates the model interpretation and identifies subset of cells that are associated to the specific phenotype label. The subset of cells with a high filter response indicates cells with a distinct signature to manifest the phenotype associated aspects of the TME. We select and perform downstream analyses for the cells with the positive filter response in most of the analysis unless stated. The relative frequency of selected cells of all input cells per patient in each tissue type as well as background were calculated and compared using a Wilcoxon rank-sum test. The best neighborhood size (*k*) of each anchor was selected when achieving by considering two parameters i) the highest classification performance (validation and test accuracy), ii) the highest frequency of selected cell significance.

We assessed the cell type composition and enrichment for selected cell subsets in *local spatial enrichment* analysis. Therefore, we defined an enrichment score (ES) that quantifies the association or exclusion of selected cells of specific cell type in the spatial proximity of the anchor cell.

We consider two quantities for calculating the ES per cell type, i) frequency of cell type with high filter response in the spatial neighborhood of the anchor cells and ii) frequency of cell type with high filter response outside of the spatial neighborhood of the anchor cells (**Fig. S25**). We define the score of cell type T to be in the nearest neighborhood of the anchor cell and selected:

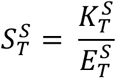

We define the score of cell type T to be in nearest neighborhood of the anchor cell:

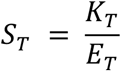

where 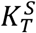 is the number of selected cell type T in the nearest neighborhood of the anchor cell, *K*_*T*_ is the number of cell type T in the nearest neighborhood of the anchor cell, *E*_*T*_ and 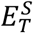 are expected value of cell type T be presented in the nearest neighborhood of the anchor cell and expected value of cell type T be selected in the nearest neighborhood of the anchor cell, respectively and calculated as follow:

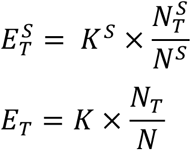

Where the variables *K* being the number of all cells in the nearest neighborhood of the anchor cell, *N* the number of all cells in the image, *N*_*T*_ the number of all cells of cell type T in the image, *K*^*S*^ the number of all selected cells in the nearest neighborhood of the anchor cell, *N*^*5*^ the number of all selected cells and 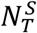 the number of all selected cells of type T.

The enrichment score for cell type T is then defined by:

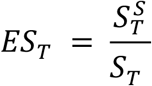

The ES values above one indicates enrichment of the selected cells of type CT in the spatial neighborhood of the anchor cell.

The enrichment score of the global spatial enrichment analysis is calculated by:

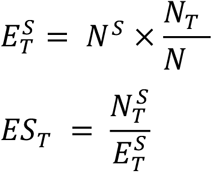

The enrichment score of a specific cell type across the patient cohort is reported as the median value of the scores across all patients. The error bars are calculated as the median absolute deviation (MAD).

### Differentially marker expression analysis

To characterize the spatially enriched cell subset, we examine if the functional marker expression in selected subsets significantly differed from non-selected one. We quantify the difference in marker expression distribution by calculating the Kolmogorov–Smirnov two-sample test statistic (KS score) for each marker per cell type. The KS score equals to 1 indicates the highest effect size between two distributions and the KS score equals to 0 indicates that two distributions are similar.

## Acknowledgments

This research was supported by the DFG Excellence Clusters EXC 2180 and EXC 2064 (SB) and the Department of Pathology and Neuropathology (CMS).

## Author contributions

Contributions according to the CRediT taxonomy where as follows: Conceptualization: M.C.; Methodology: S.B., M.C.; Software: S.B.; Validation S.B., J.C., M.C., C.M.S., A.M., M.Z; Formal analysis: S.B., J. C., M.C.; Investigation: S.B., M.C., C.M.S.; Resources S.B; Data Curation: S.B; Writing – original draft preparation: S.B. M.C.; Writing – review and editing: M.C., S.B., C.M.S., K.W.H.; Visualization: S.B., M.C., C.M.S., A.M., M.Z; Supervision: M.C., C.M.S. All authors edited the manuscript and approved its final version.

## Data and materials availability

The code of the S^3^-CIMA workflow has been deposited at https://github.com/claassenlab/S3-CIMA.

