## Supplementary Figures for "S^3^-CIMA: Supervised spatial single-cell image analysis for the identification of disease-associated cell type compositions in tissue"

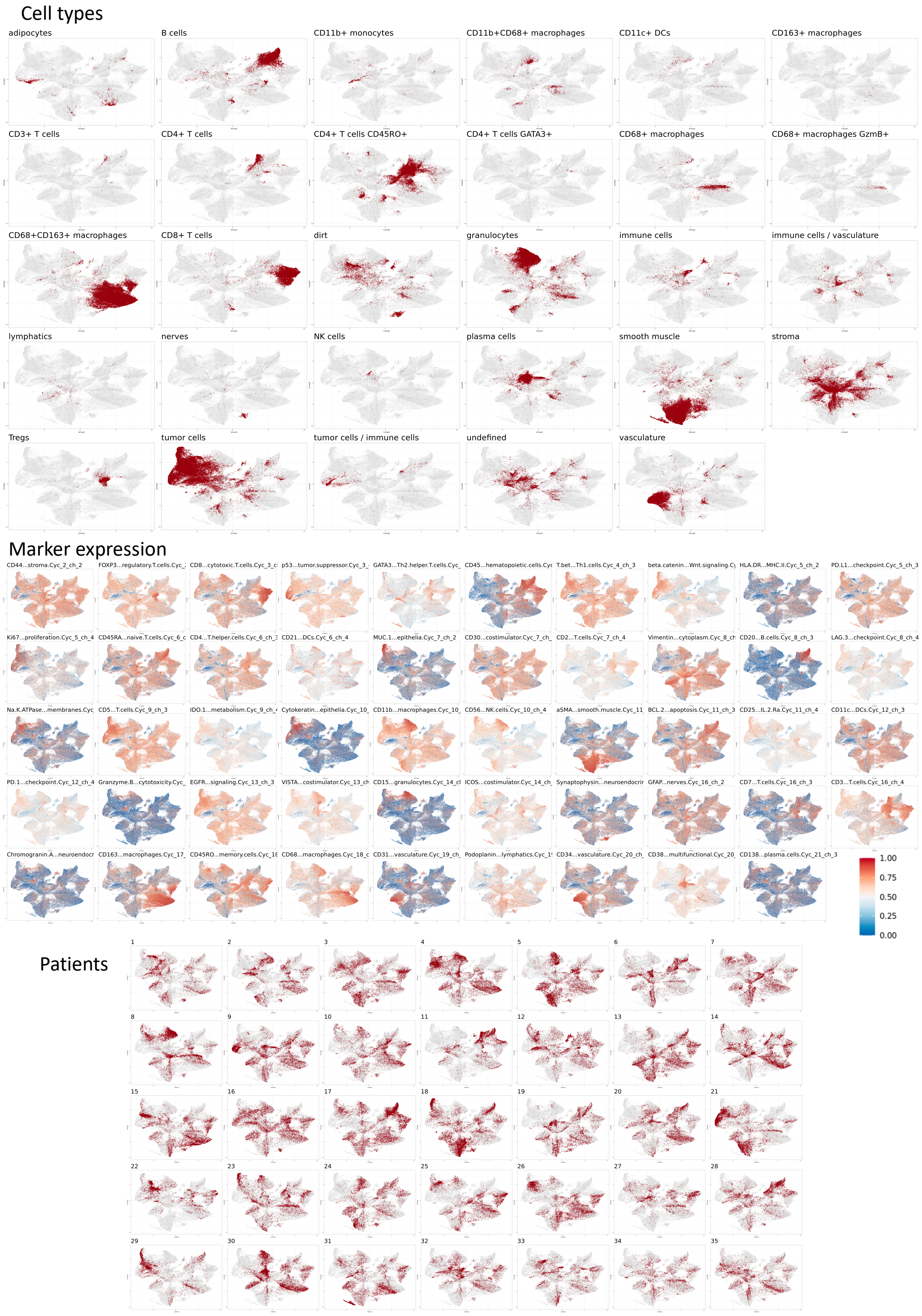


**Fig. S1.**

CRC data

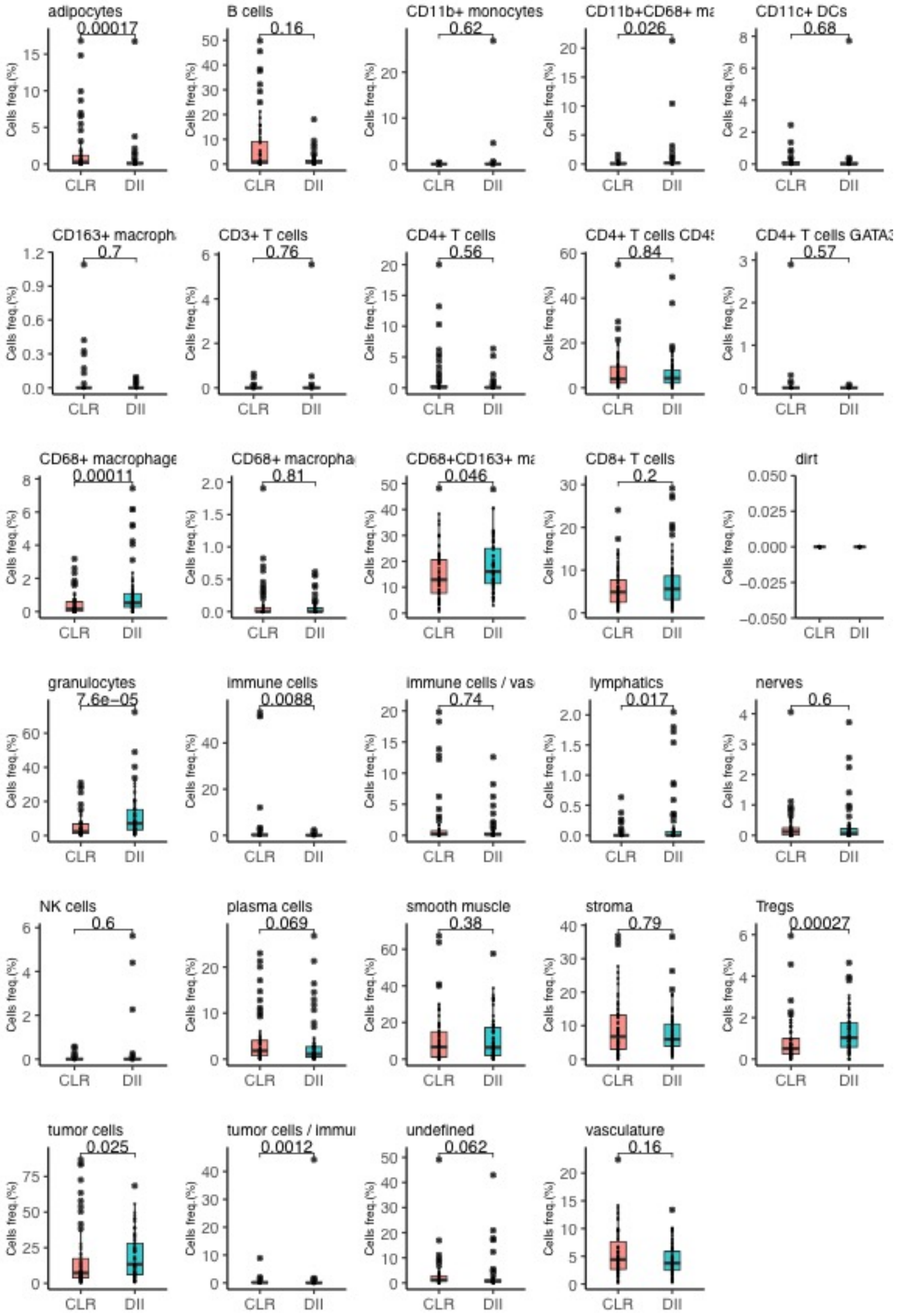


**Fig. S2.**

Box plot of frequency of each cell type in each CRC group (CLR vs DII)

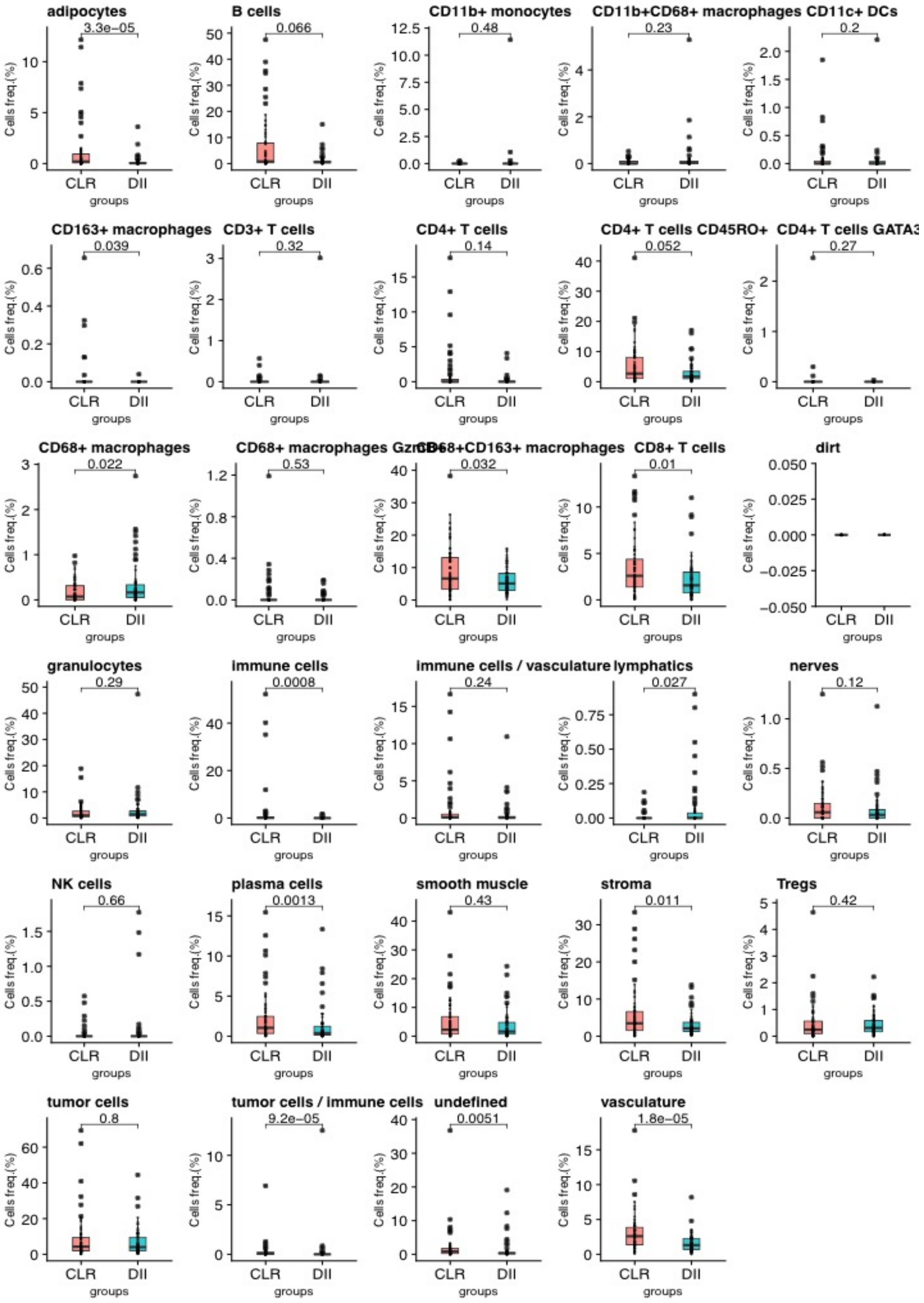


**Fig. S3.**

Box plot of frequency of selected cell obtained from $S^3$-CIMA global enrichment analysis at k = 30 in each CRC group per cell type.

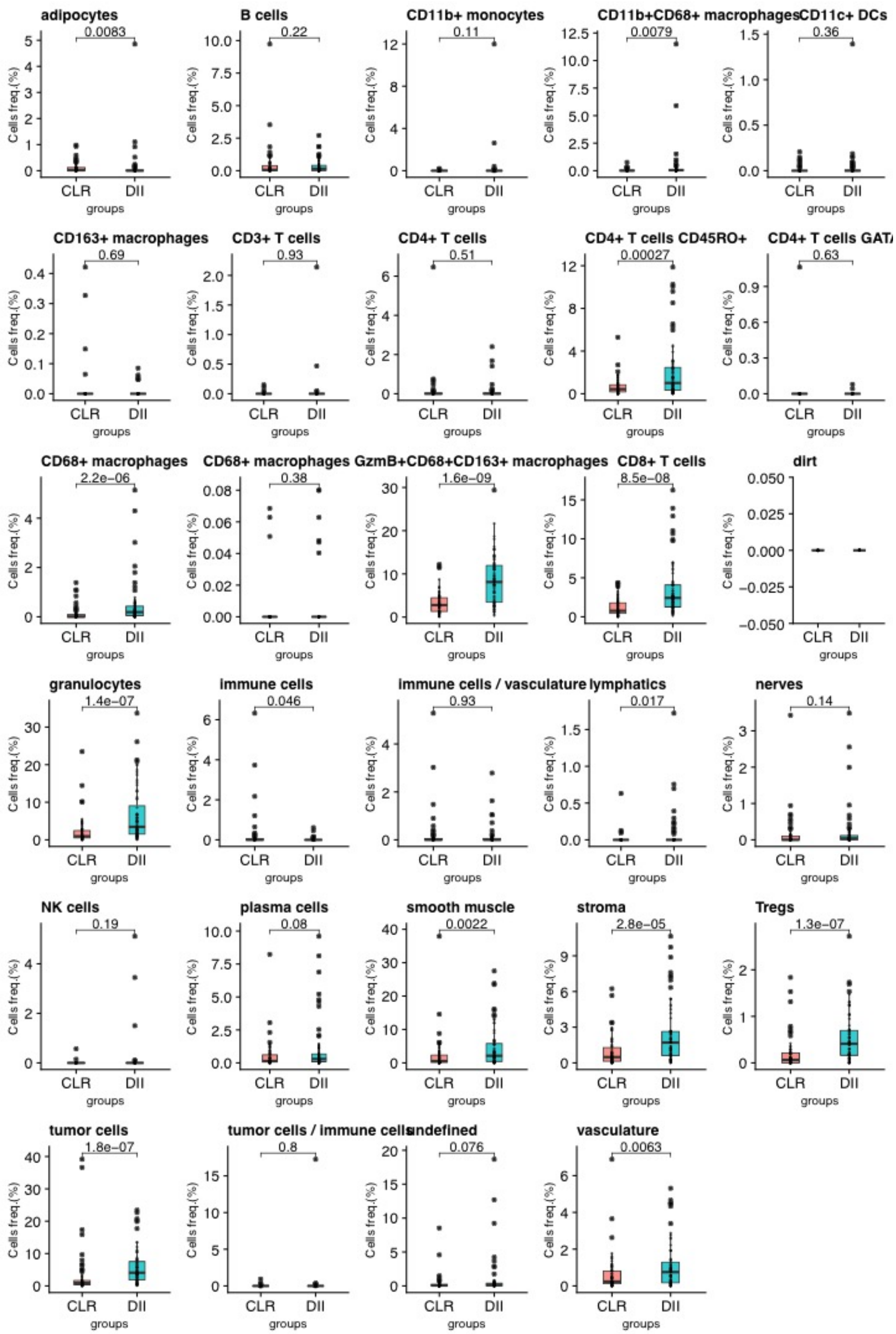


**Fig. S4.**

Box plot of frequency of selected cell obtained from $S^3$-CIMA global enrichment analysis at k = 50 in each CRC group per cell type.

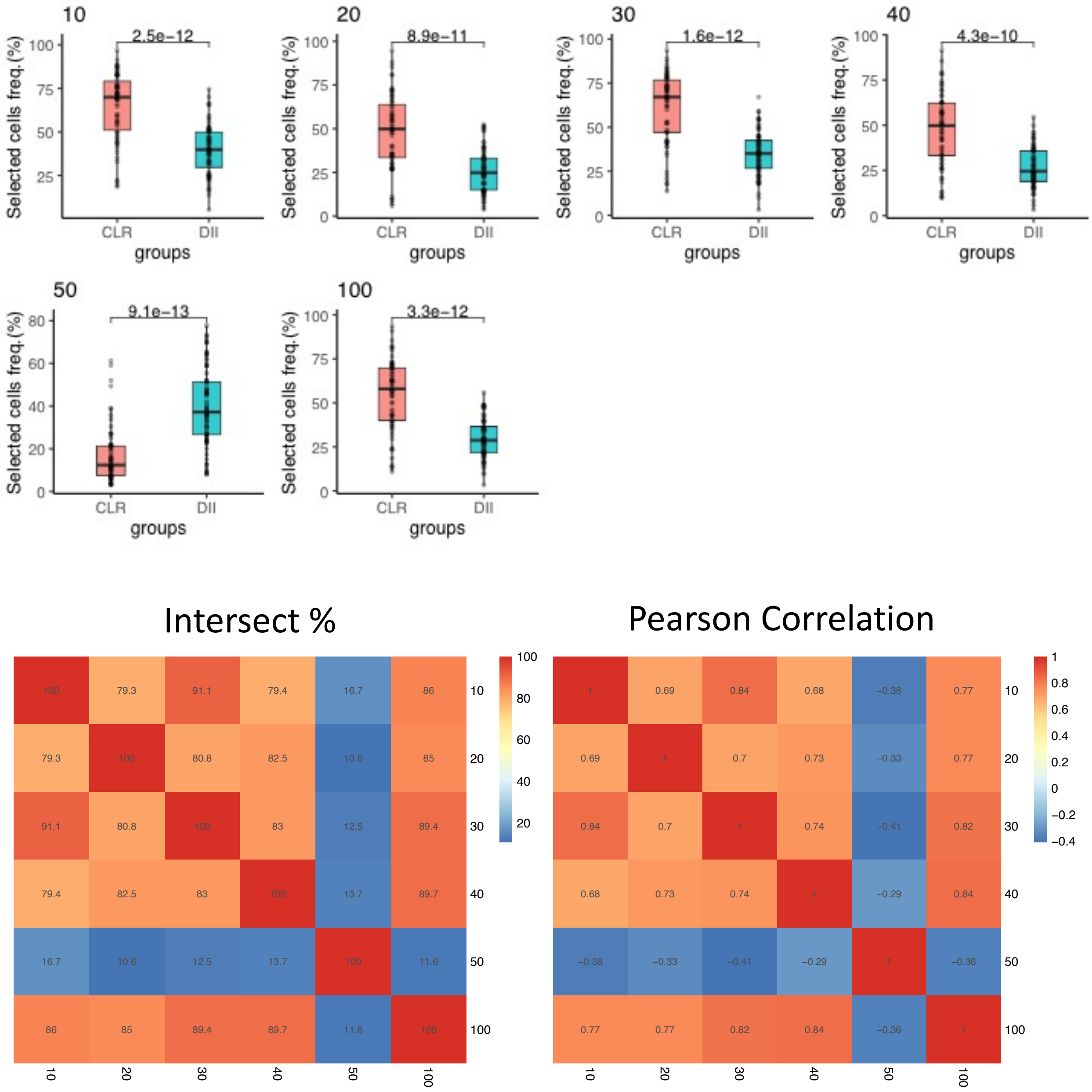


**Fig. S5.**
The frequency of selected cells between groups using $S^3$-CIMA global enrichment analysis across different cell neighborhood sizes (10 to 100). Heatmaps show the overlap (%) and correlation between the set of selected cells over the different neighborhood sizes.

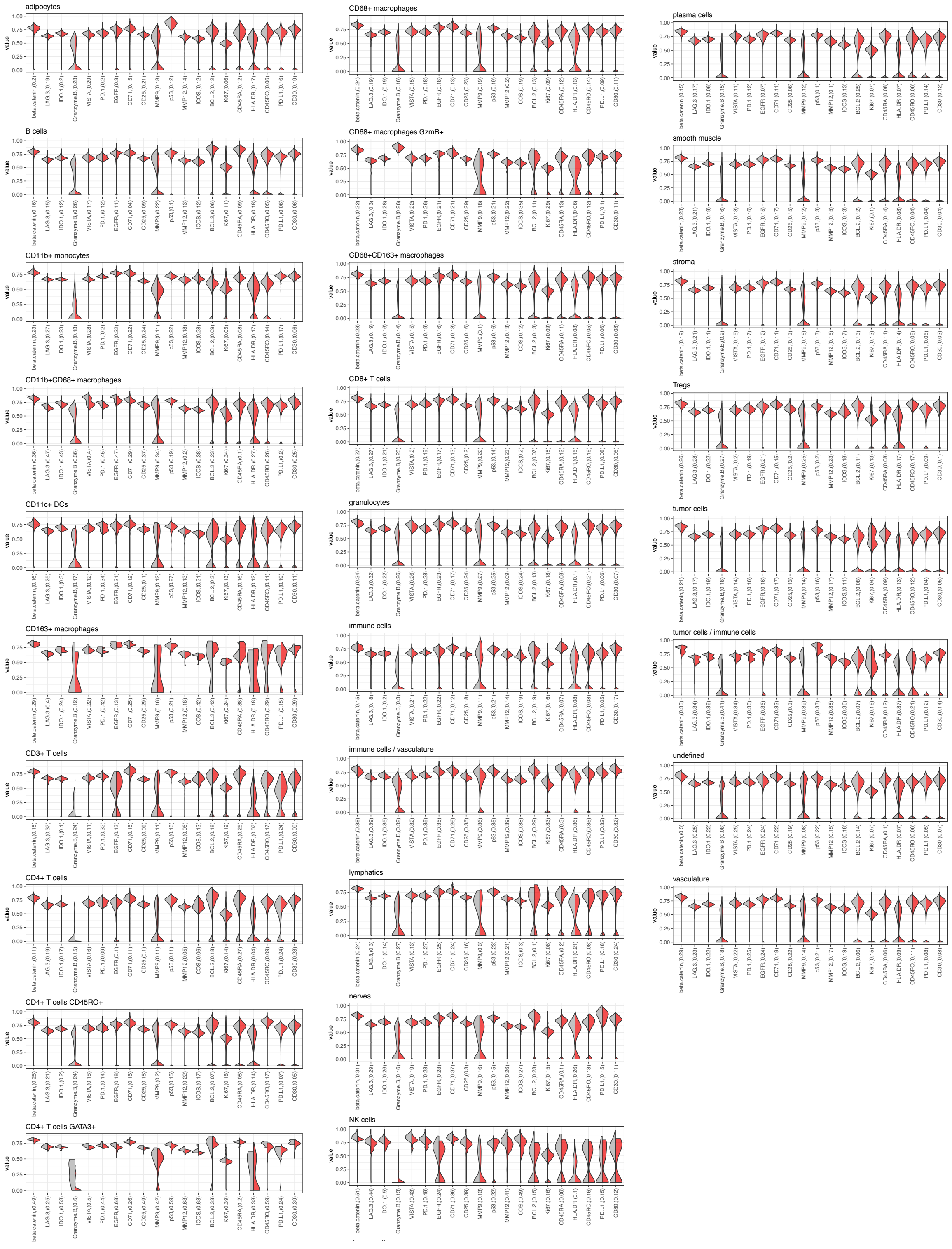


**Fig. S6.**

S$^3$-CIMA global enrichment analysis at k =30, Density of functional marker expression showing greatest differential abundance in terms of the Kolmogorov–Smirnov two-sample test between the selected and non-selected cell subsets per cell types.

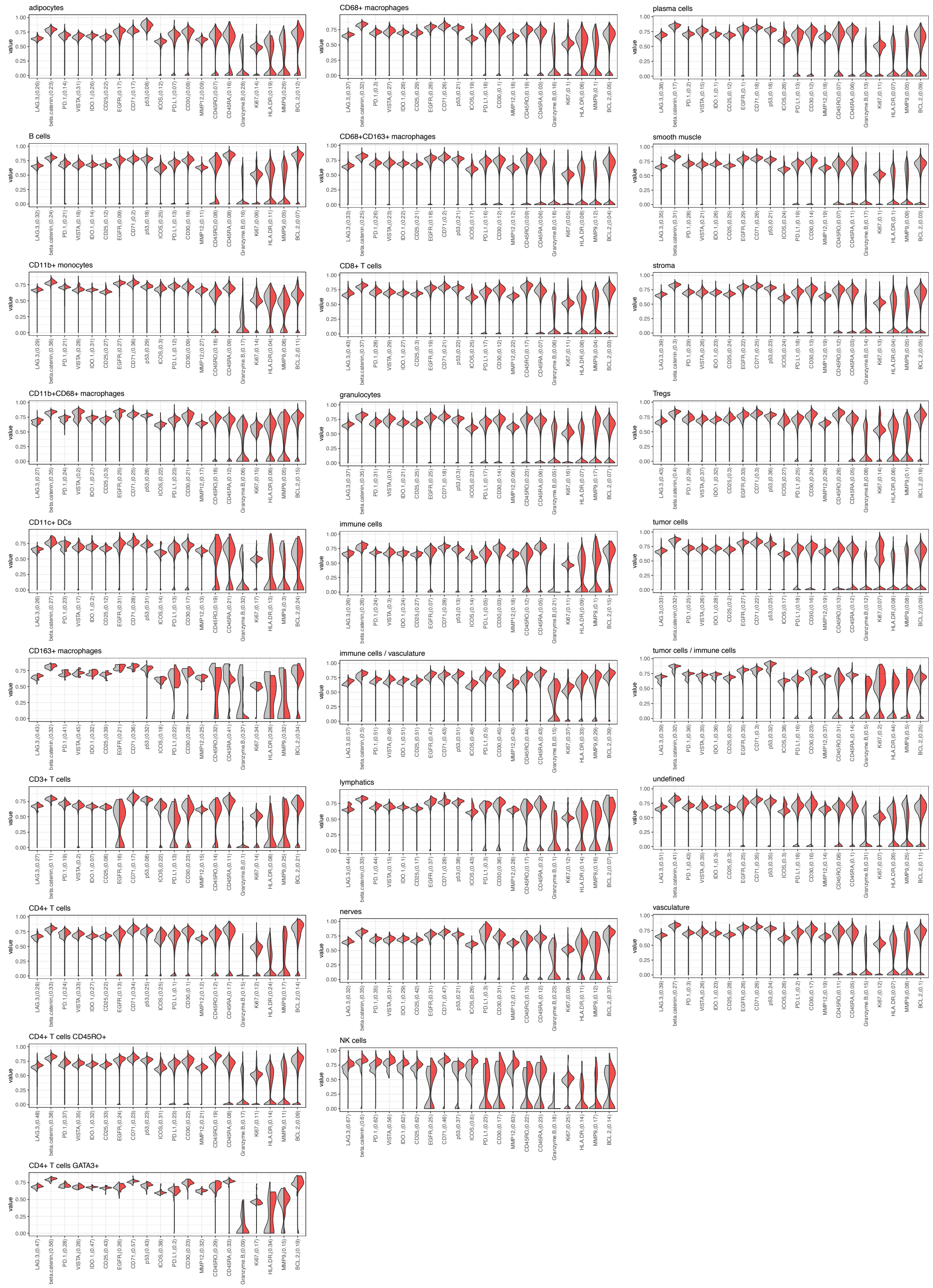


**Fig. S7.**

$S^3$-CIMA global enrichment analysis at k =50, Density of functional marker expression showing greatest differential abundance in terms of the Kolmogorov–Smirnov two-sample test between the selected and non-selected cell subsets per cell types.

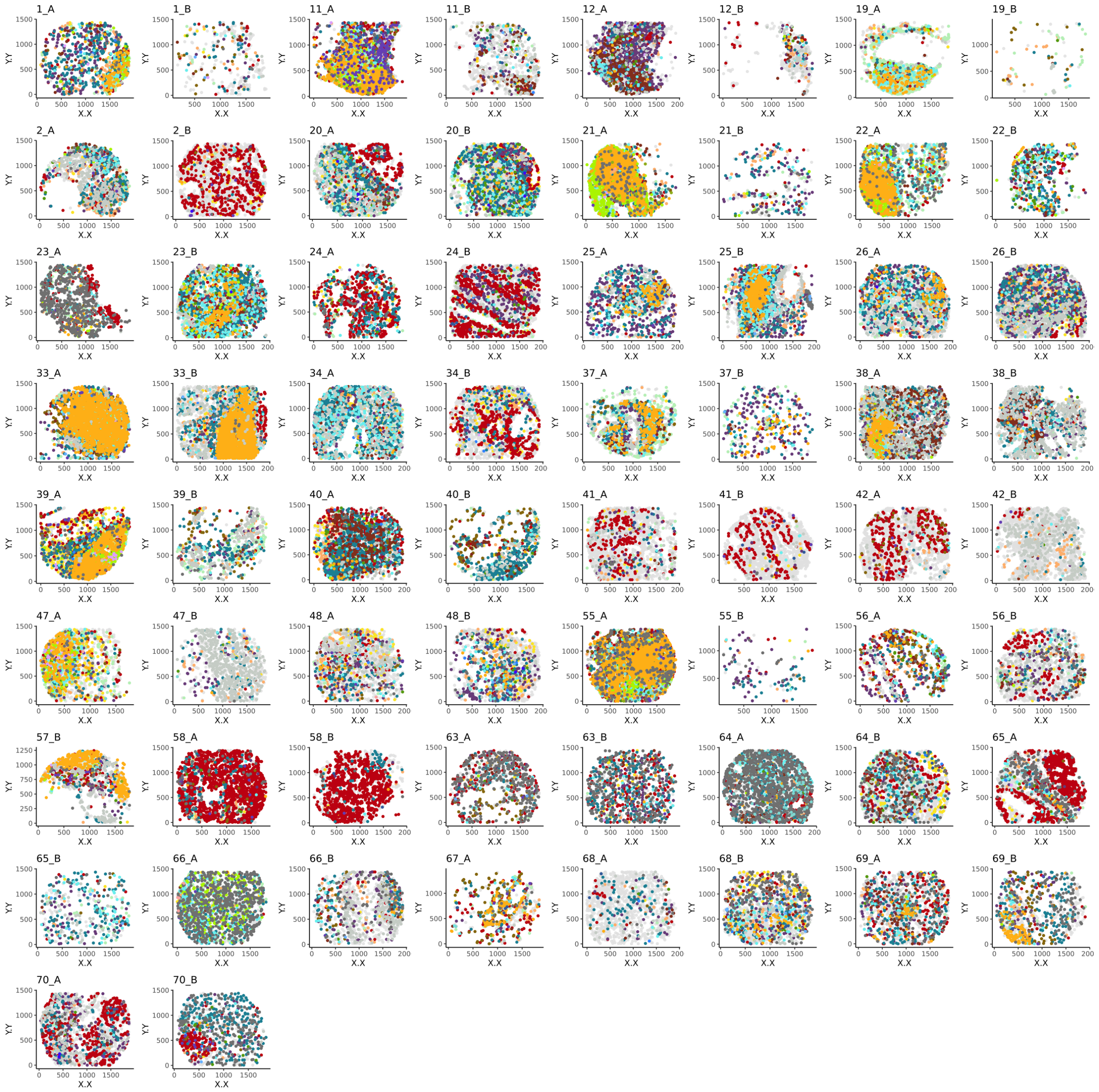


**Fig. S8.**

S³-CIMA global enrichment analysis at k =30, Selected cells (colored by cell type) are mapped back to the corresponding patient CODEX images in both CLR group.

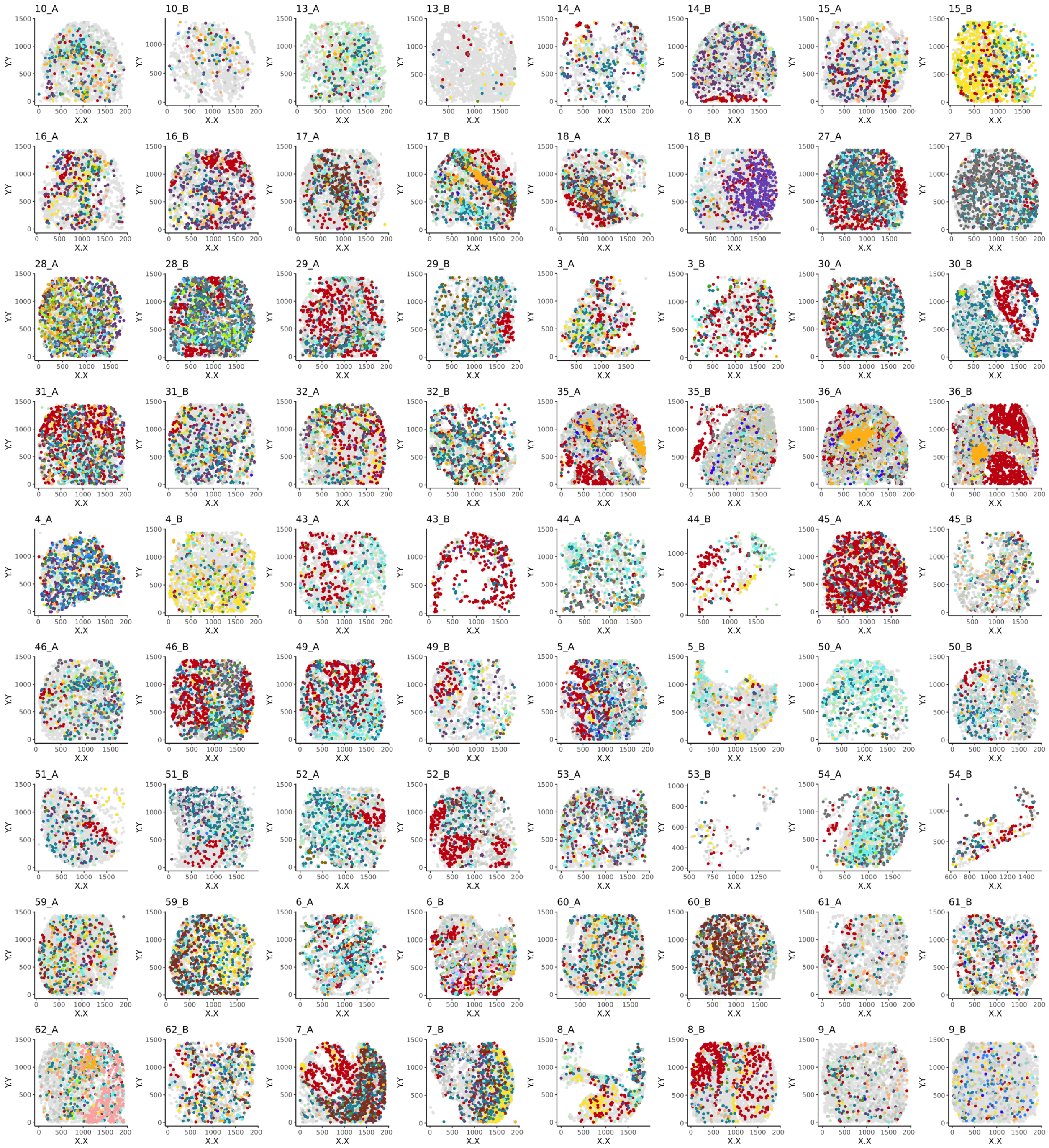


**Fig. S9.**

S[3]-CIMA global enrichment analysis at k =30, Selected cells (colored by cell type) are mapped back to the corresponding patient CODEX images in both DII group.

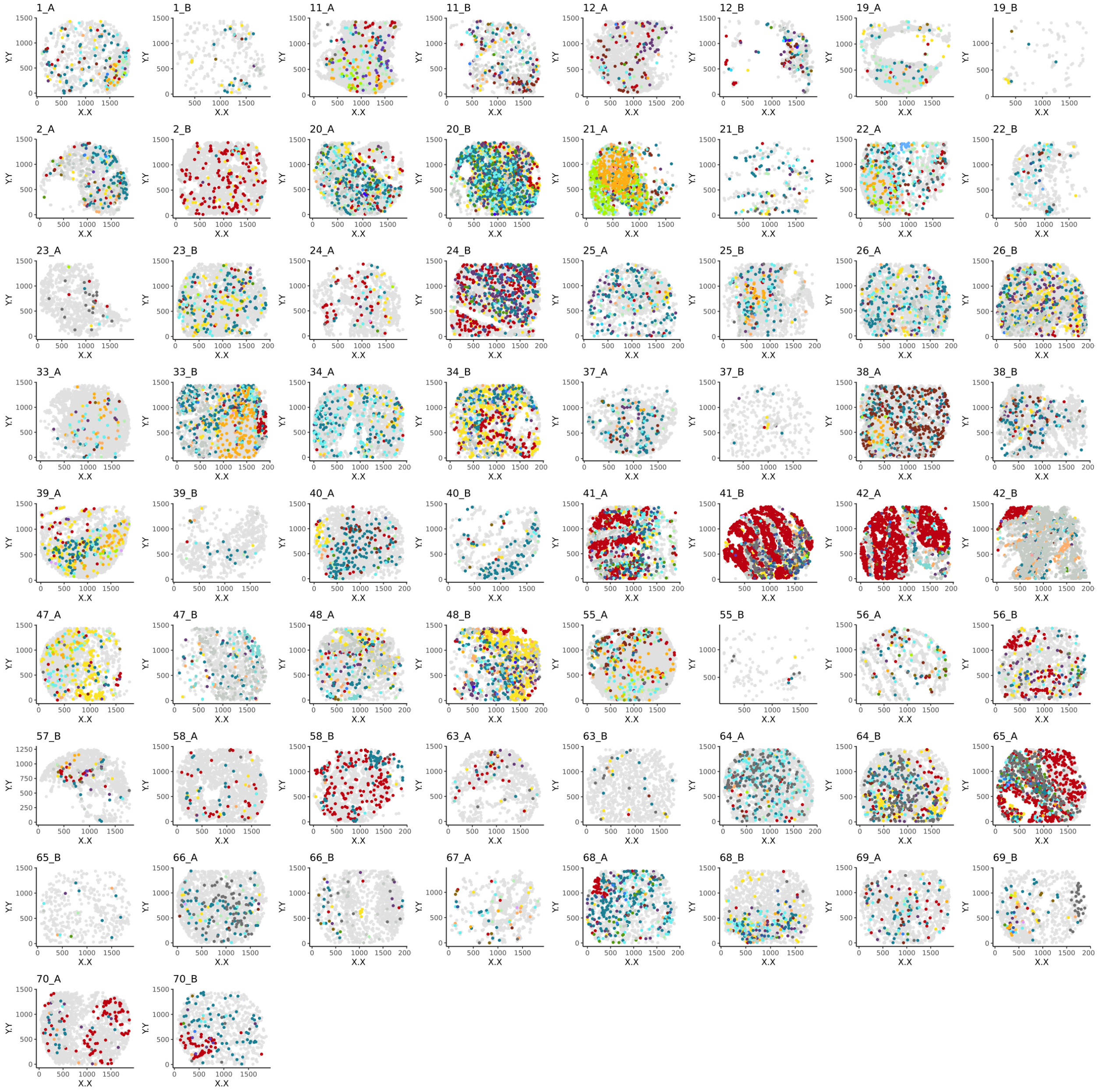


**Fig. S10.**

S$^3$-CIMA global enrichment analysis at k =50, Selected cells (colored by cell type) are mapped back to the corresponding patient CODEX images in both CLR group.

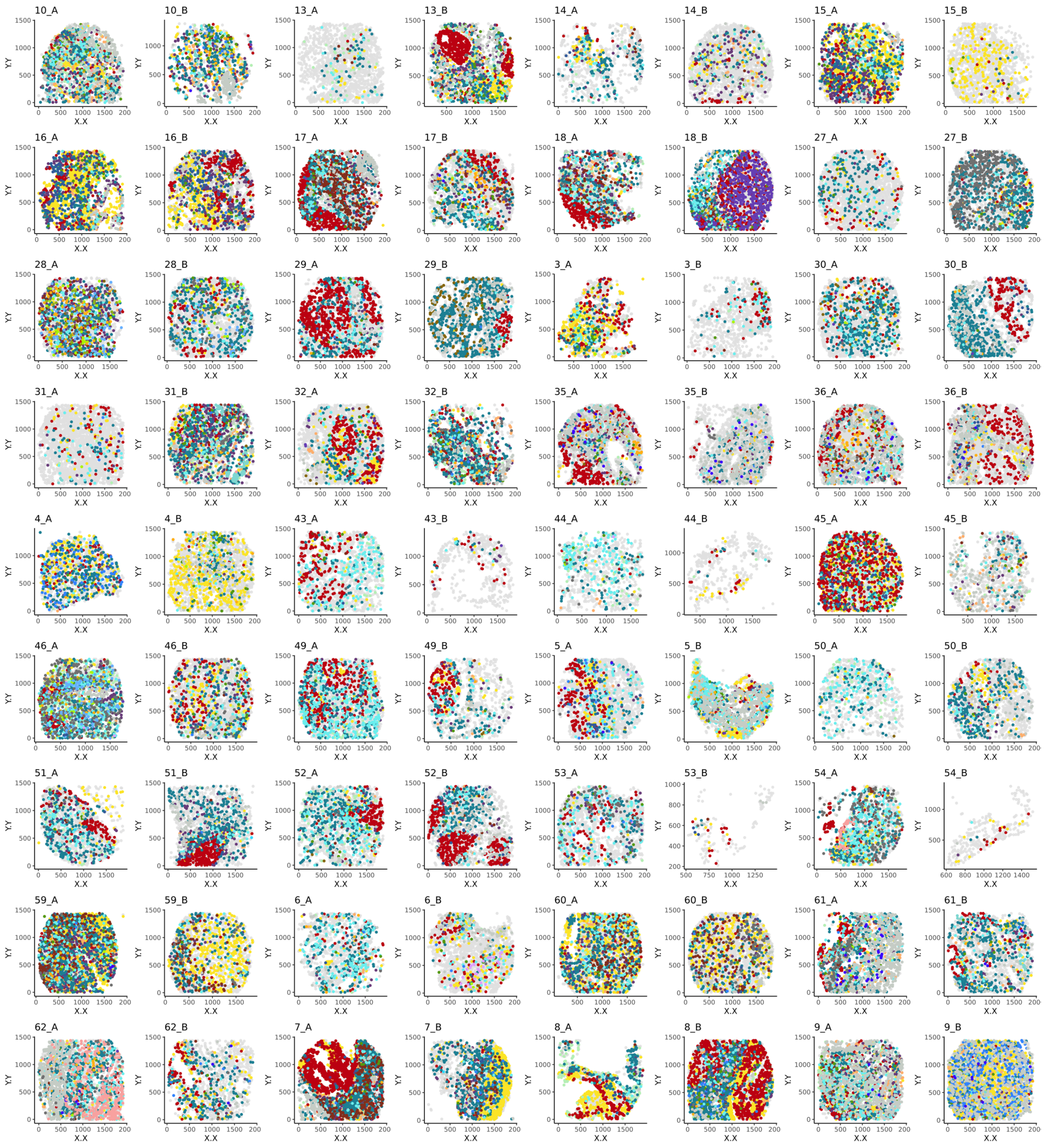


**Fig. S11.**

$S^3$-CIMA global enrichment analysis at k =50, Selected cells (colored by cell type) are mapped back to the corresponding patient CODEX images in both DII group.

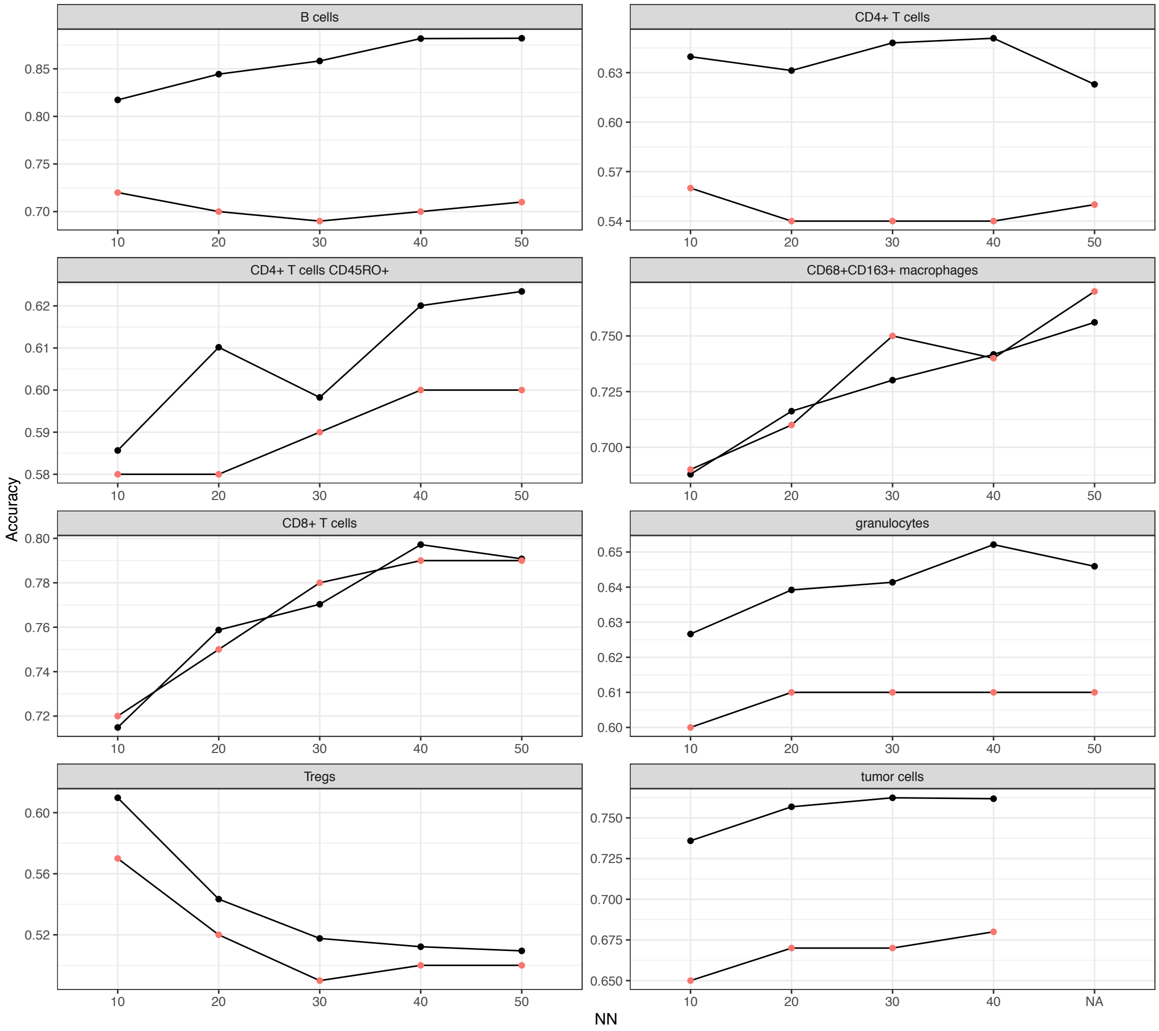


**Fig. S12.**

S[3]-CIMA anchor based spatial enrichment analysis classification performance (test (red) and train (black) accuracy) across different cell neighborhood sizes (10 to 50).

Anchor: granulocyte

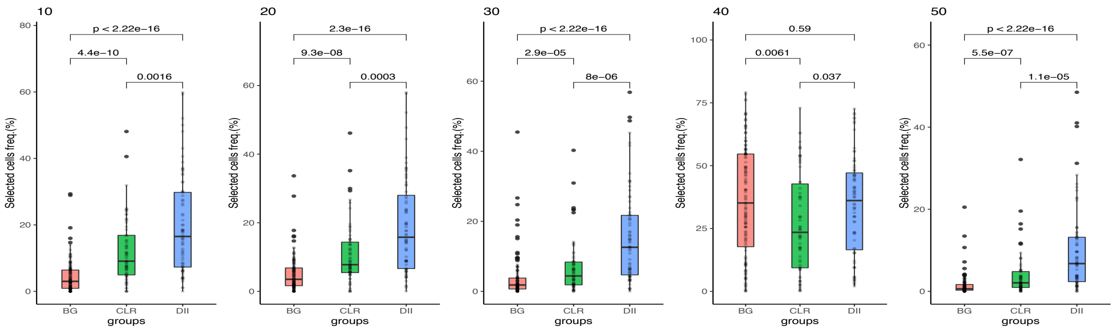


Anchor: CD4+ T cells CD45RO+

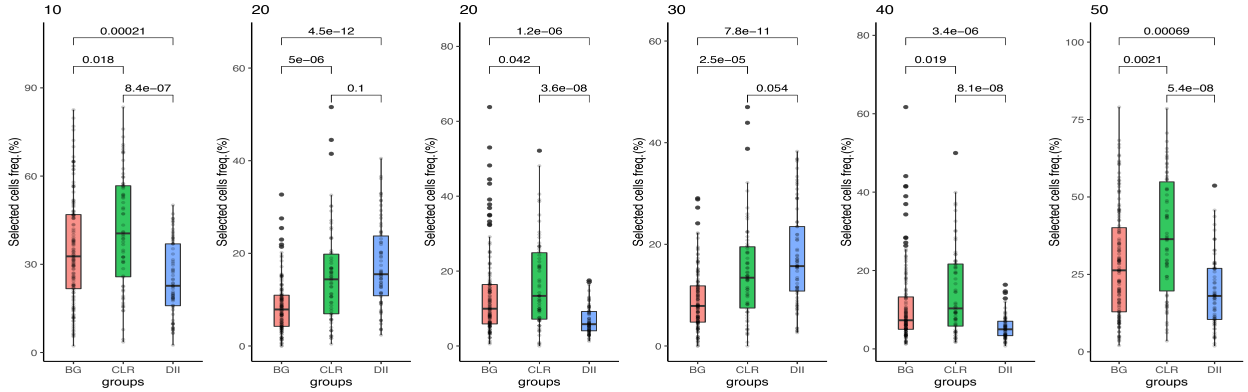


**Fig. S13.**

Boxplots of selected cell type frequency of two CRC groups using $S^3$-CIMA anchor based spatial enrichment analysis with granulocyte and CD4+ T cells CD45RO+ as the anchor.

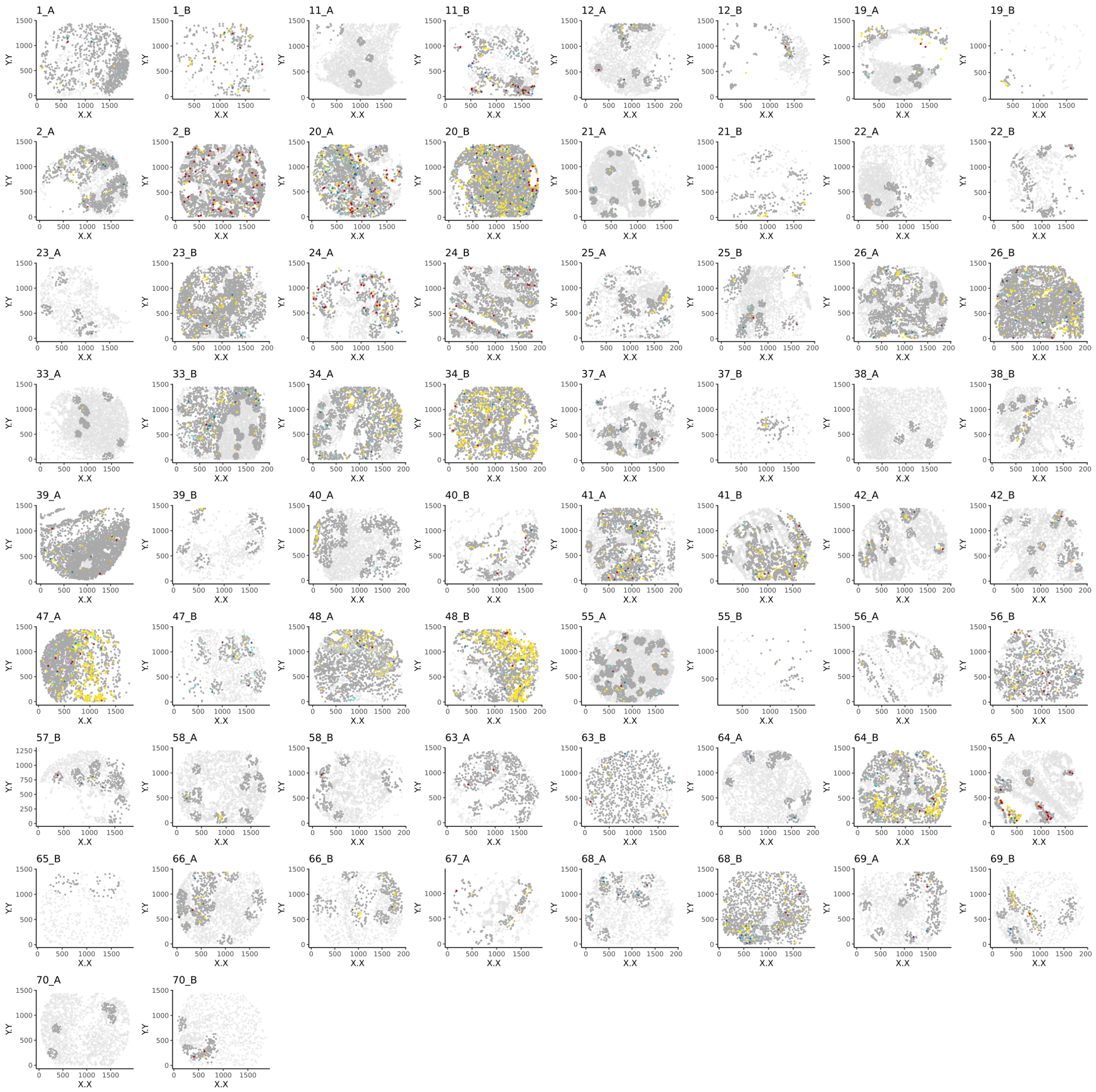


**Fig. S14.**

Selected cells (colored by cell type) using $S^3$-CIMA anchor based spatial enrichment analysis with granulocyte as the anchor mapped back to the corresponding patient CODEX images in the **CLR** group.

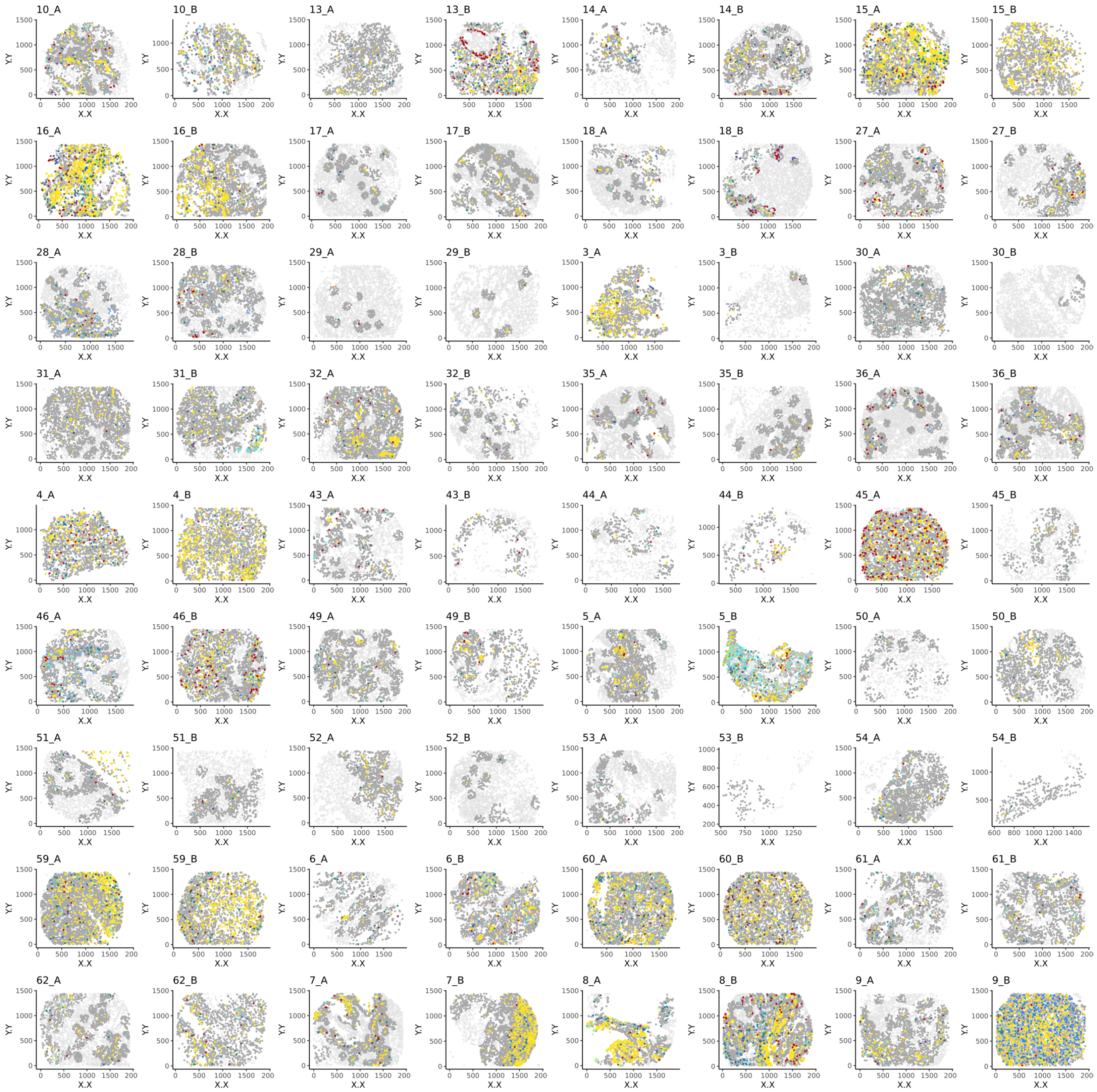


**Fig. S15.**

Selected cells (colored by cell type) using S$^3$-CIMA anchor based spatial enrichment analysis with granulocyte as the anchor mapped back to the corresponding patient CODEX images in the **DII** group.

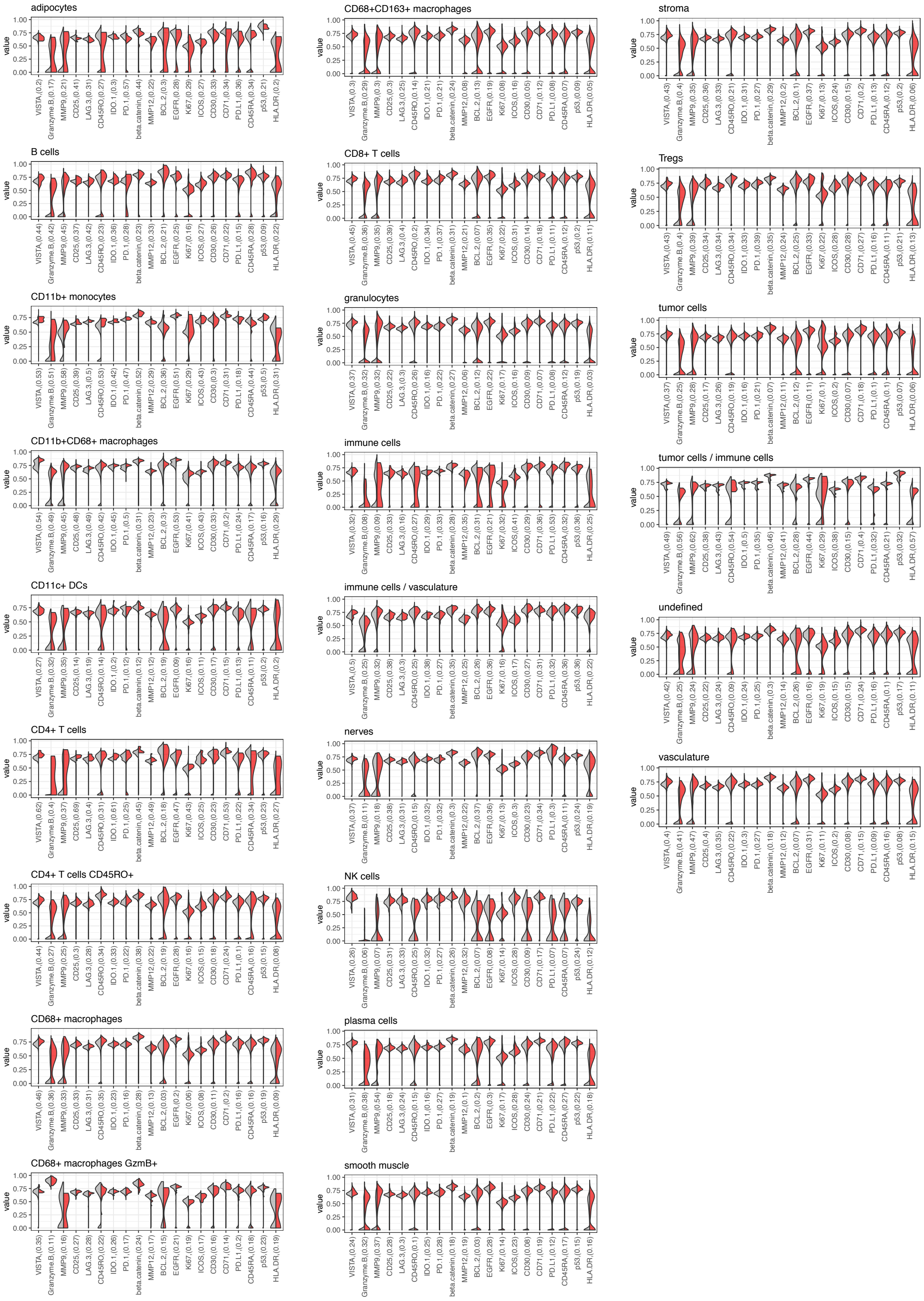


**Fig. S16.**

S[3]-CIMA local enrichment analysis at k =30 granulocyte as the anchor, Density of functional marker expression showing differential abundance (KS two-sample test) between the selected and non-selected cells per cell types.

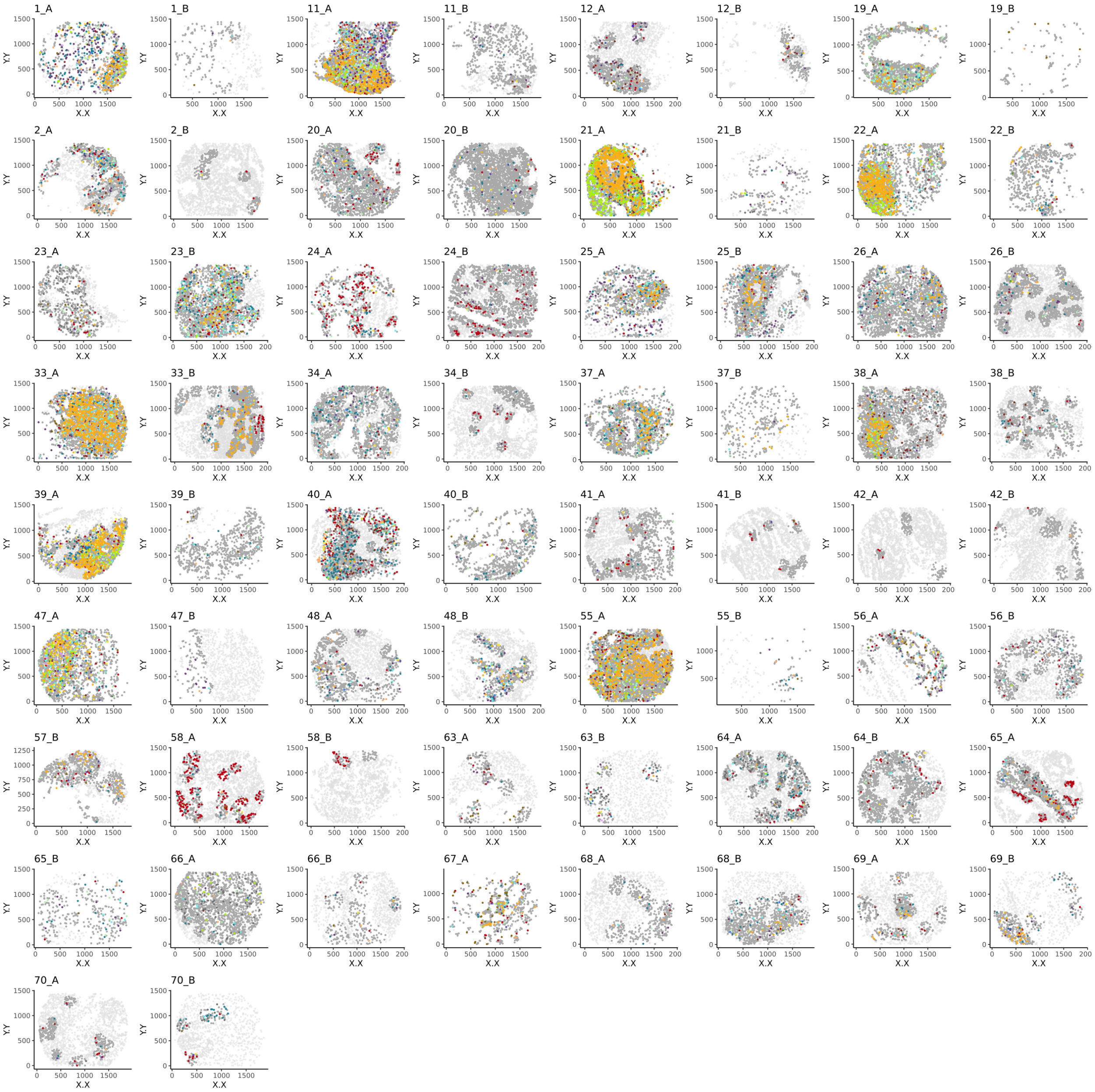


**Fig. S17.**

Selected cells (colored by cell type) using S$^3$-CIMA anchor based spatial enrichment analysis with CD4+ T cells CD45RO+ as the anchor mapped back to the corresponding patient CODEX images in the **CLR** group.

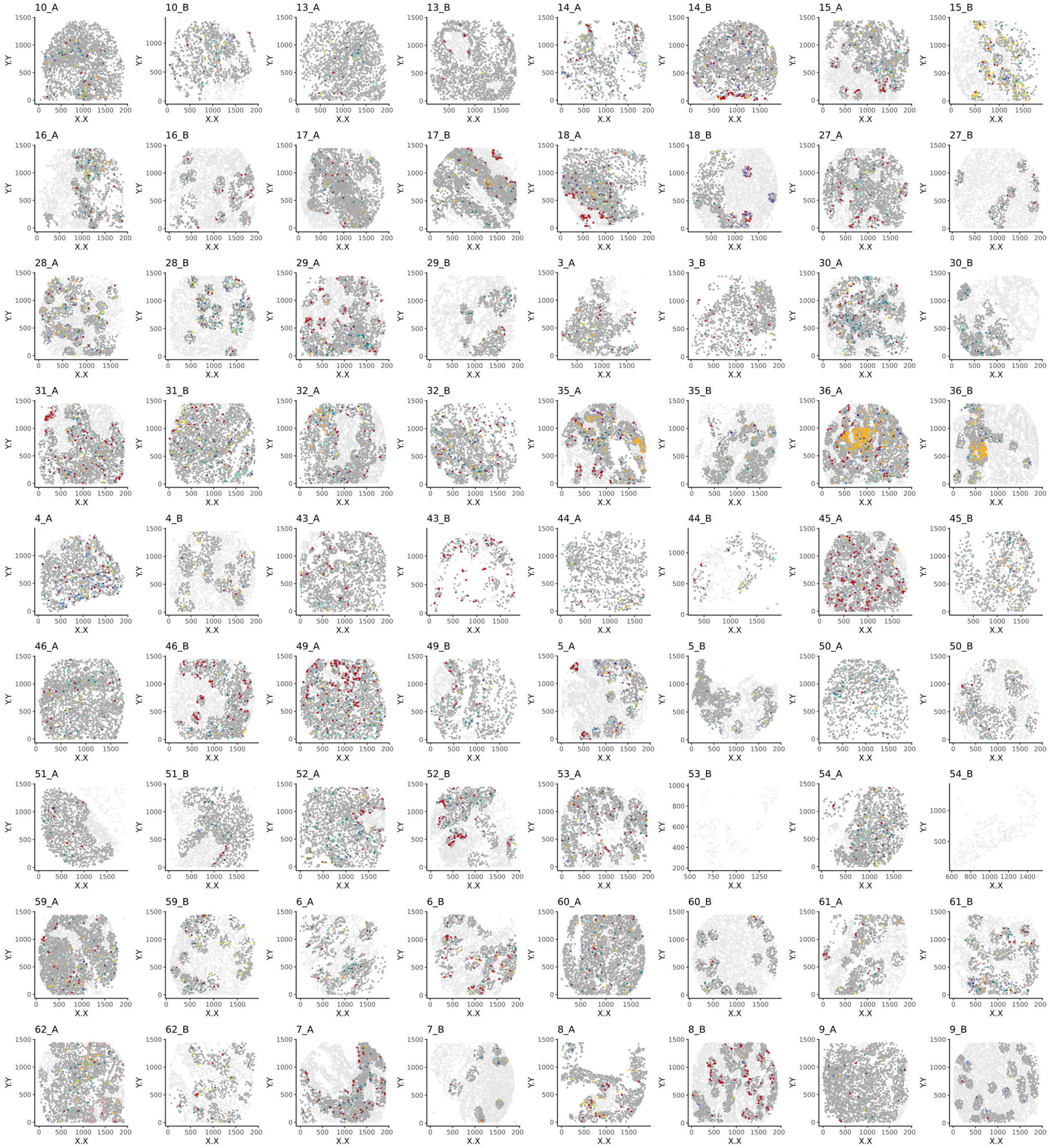


**Fig. S18.**

Selected cells (colored by cell type) using $S^3$-CIMA anchor based spatial enrichment analysis with CD4+ T cells CD45RO+ as the anchor mapped back to the corresponding patient CODEX images in the **DII** group.

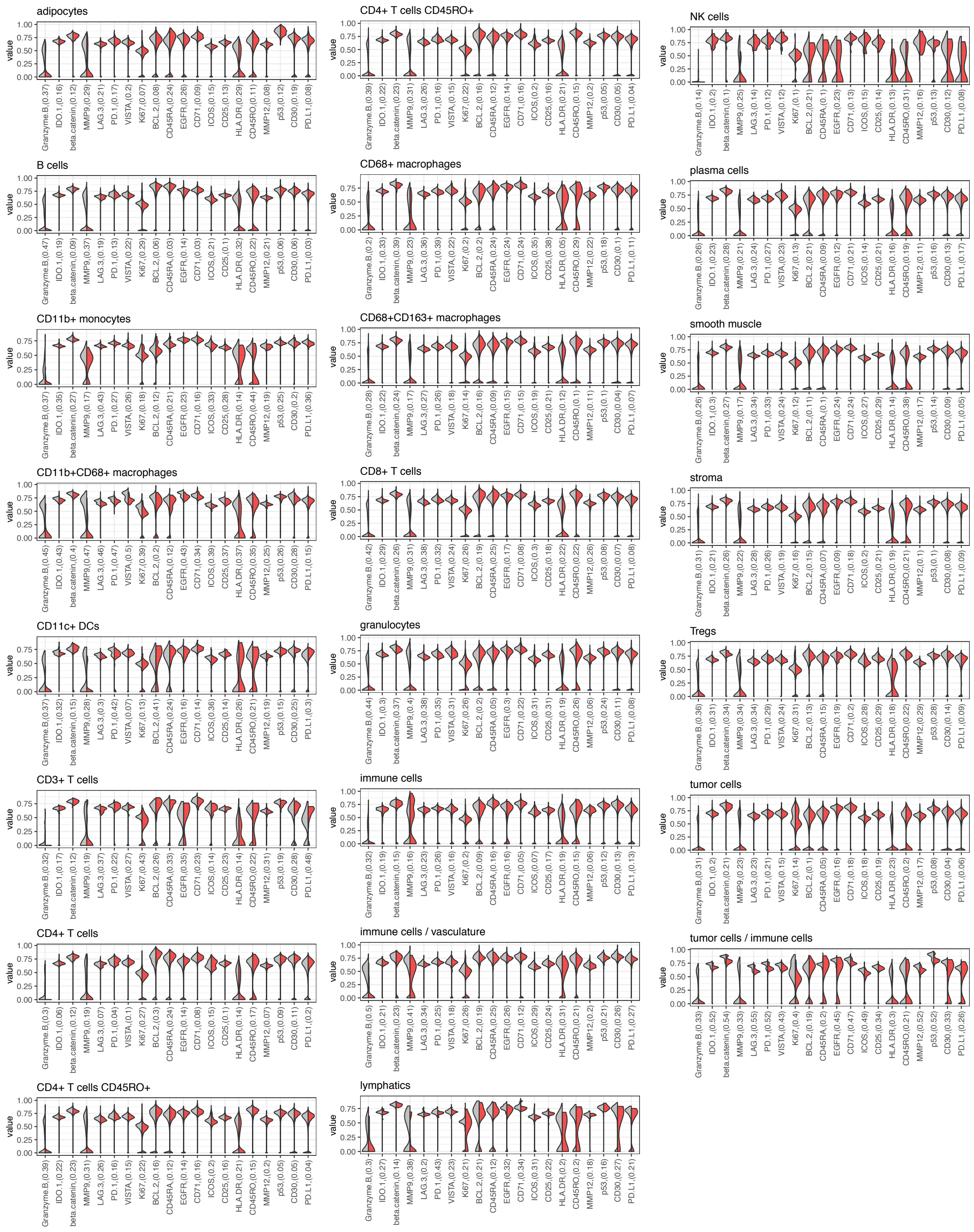


**Fig. S19.**

$S^3$-CIMA local enrichment analysis at k =40, CD4+ T cells CD45RO+ as the anchor, Density of functional marker expression showing differential abundance (KS two-sample test) between the selected and non-selected cells per cell types.

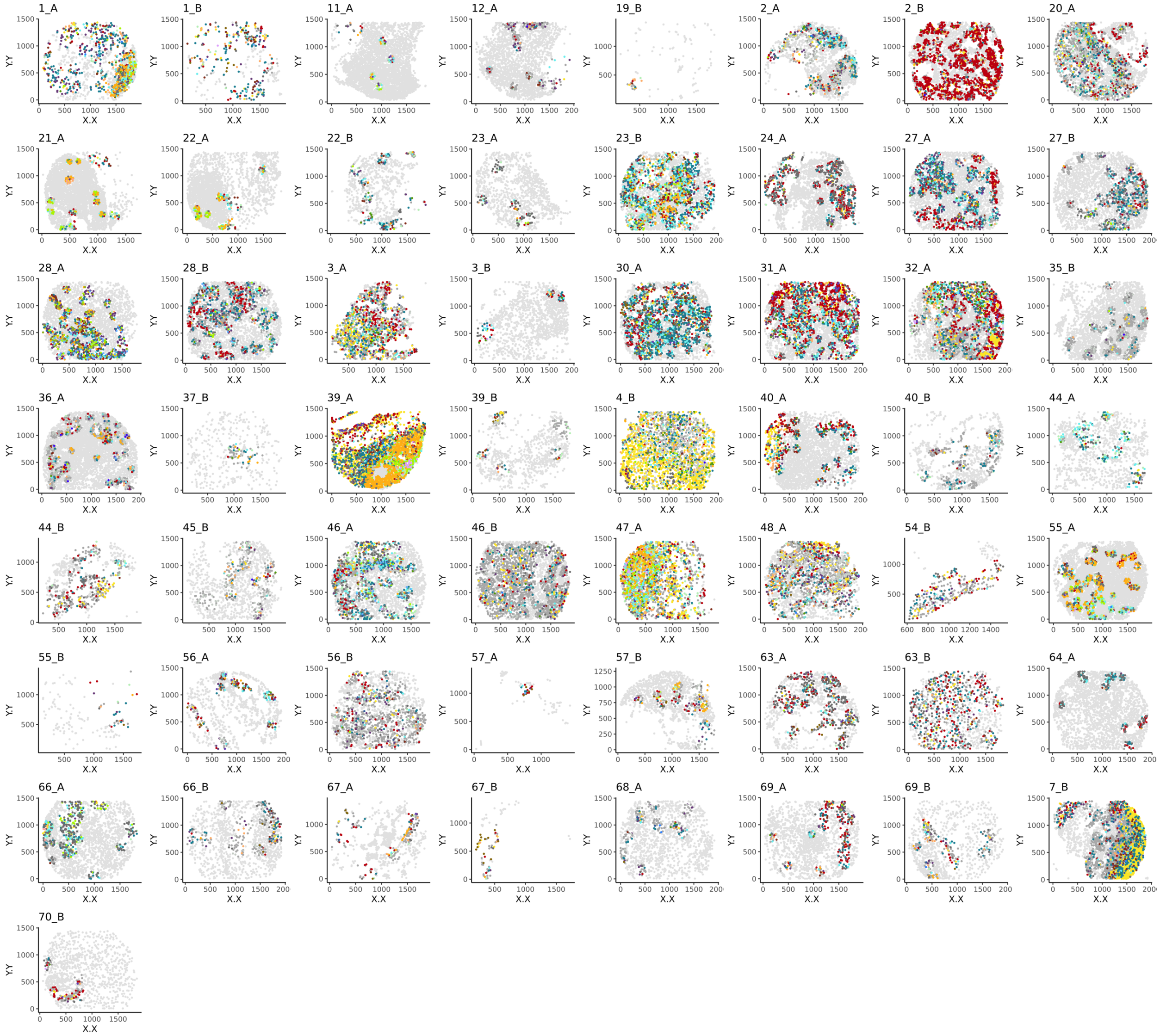


**Fig. S20.**

Selected cells (colored by cell type) using S$^3$-CIMA functional spatial enrichment analysis with granulocytes as the anchor and EGFR marker expression as the phenotype mapped back to the corresponding patient CODEX images in the **EGFR low** group. (Filter 1).

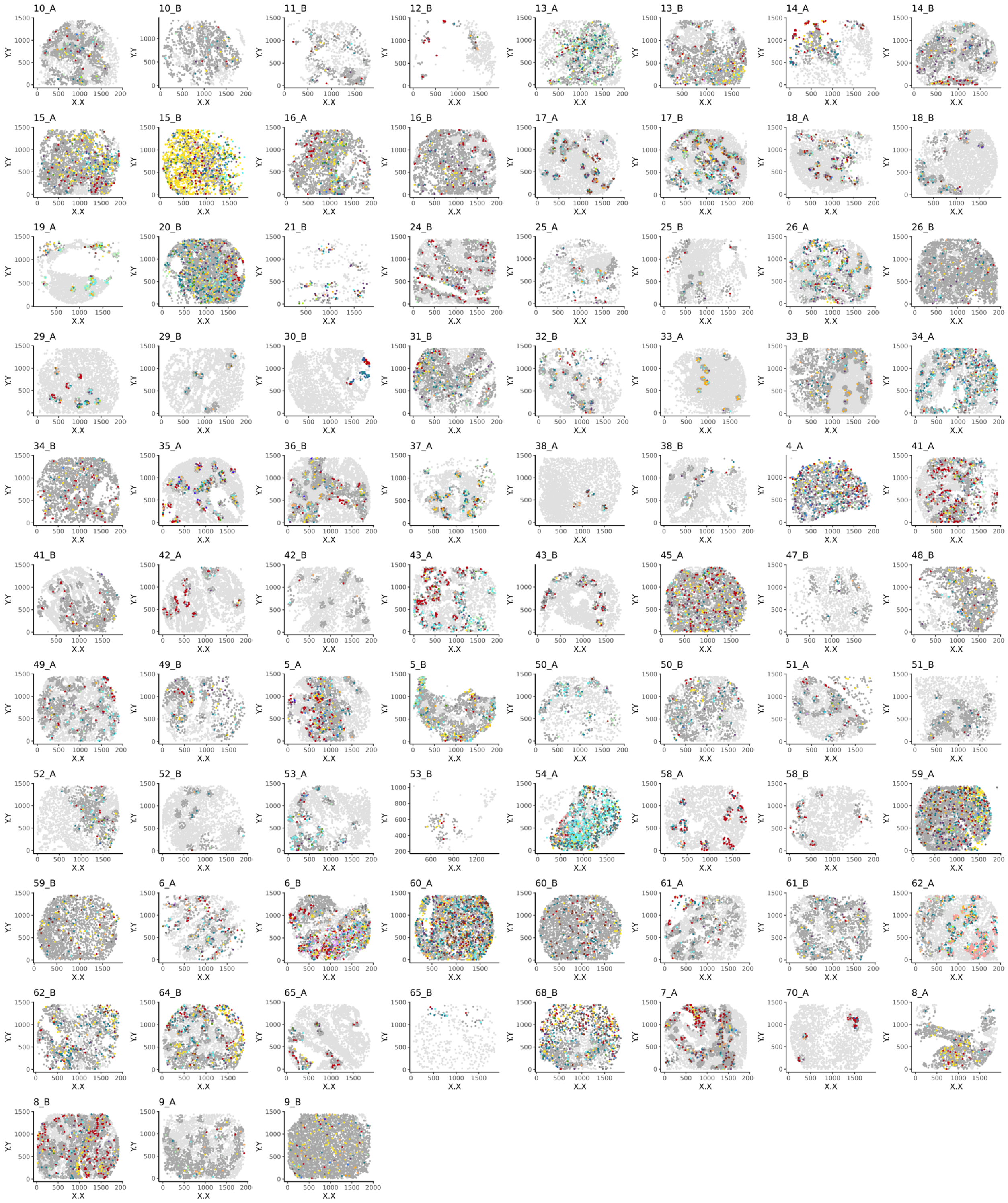


**Fig. S21.**

Selected cells (colored by cell type) using $S^3$-CIMA functional spatial enrichment analysis with granulocytes as the anchor and EGFR marker expression as the phenotype mapped back to the corresponding patient CODEX images in the **EGFR high** group. (Filter 1).

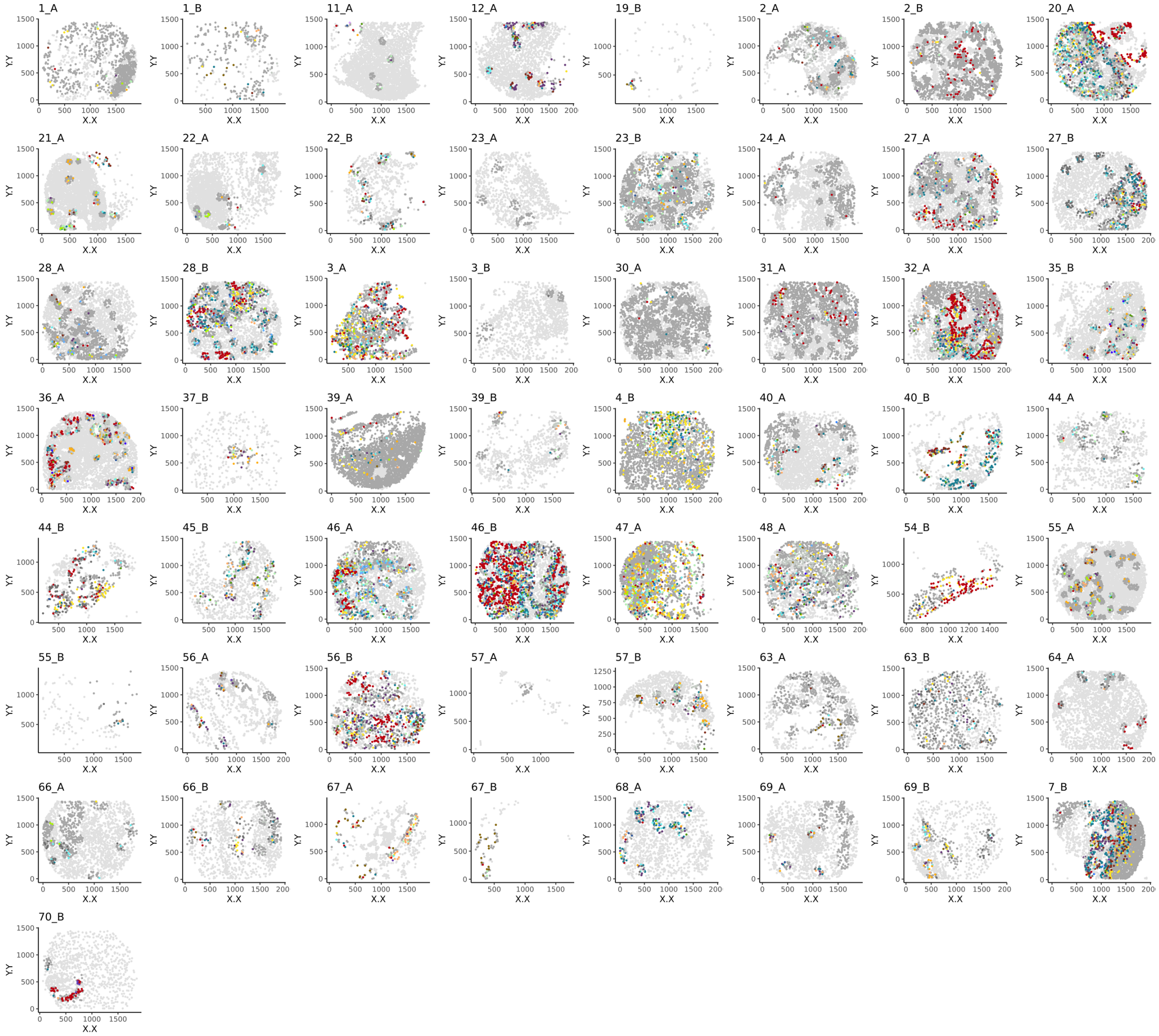


**Fig. S22.**

Selected cells (colored by cell type) using $S^3$-CIMA functional spatial enrichment analysis with granulocytes as the anchor and EGFR marker expression as the phenotype mapped back to the corresponding patient CODEX images in the **EGFR low** group. (Filter 2) .

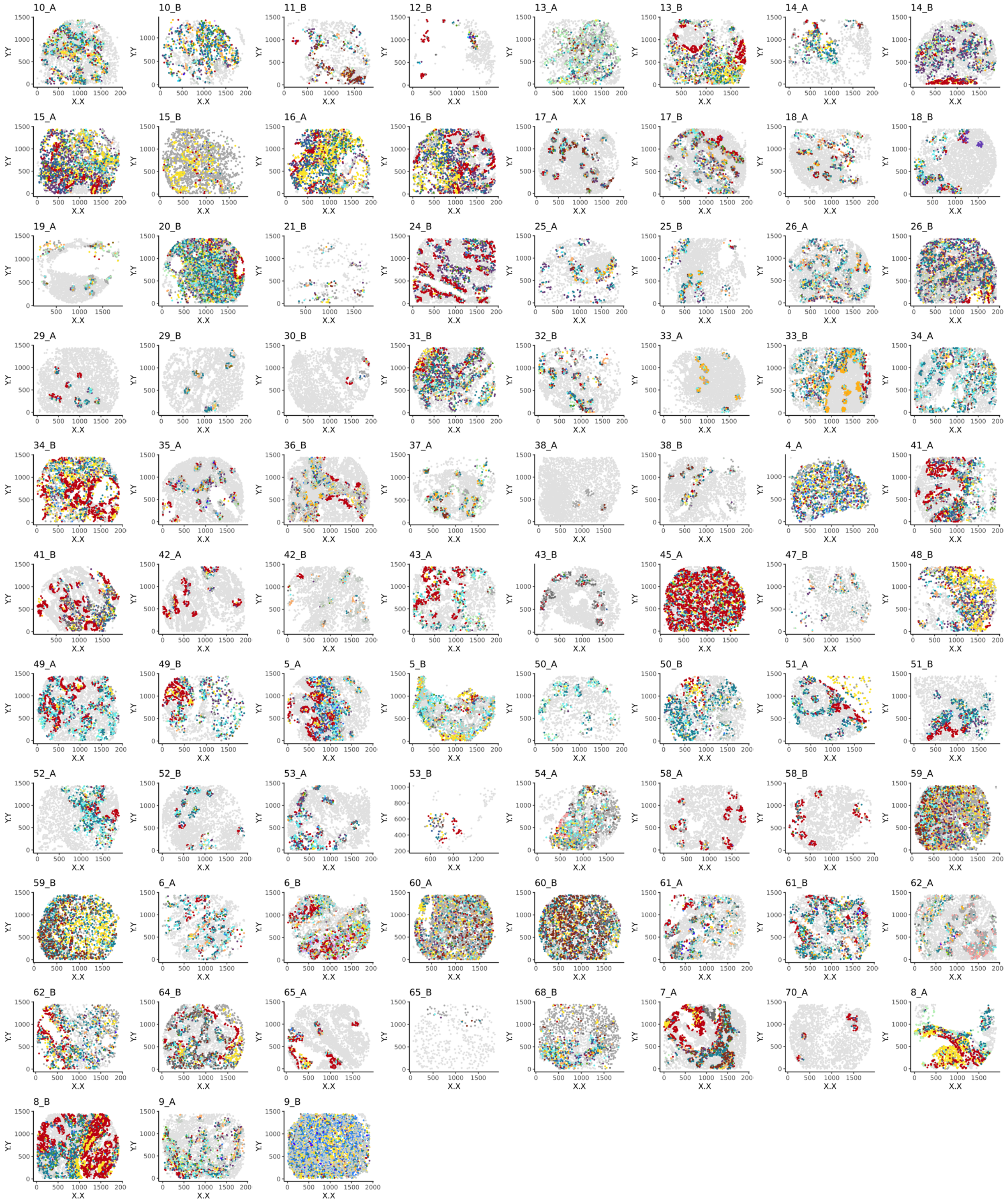


**Fig. S23.**

Selected cells (colored by cell type) using S[3]-CIMA functional spatial enrichment analysis with granulocytes as the anchor and EGFR marker expression as the phenotype mapped back to the corresponding patient CODEX images in the **EGFR high** group. (Filter 2).

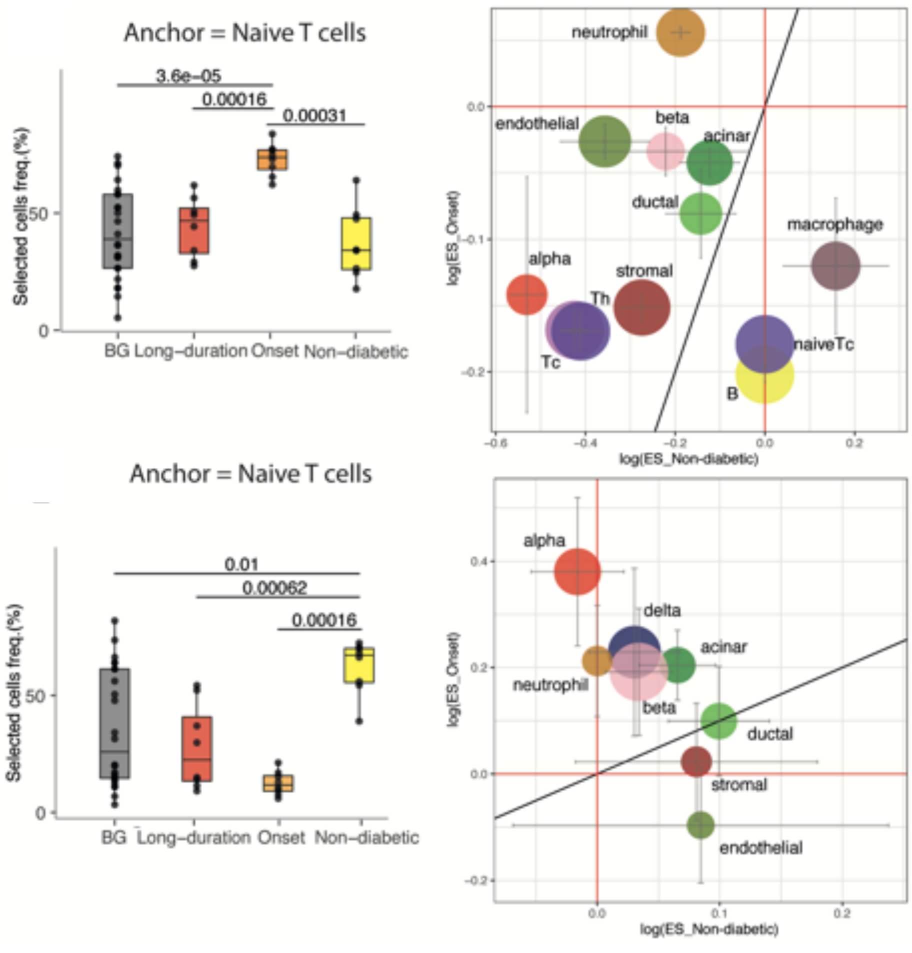


**Fig. S24.**
S[3]-CIMA anchor based spatial enrichment analysis with Naïve T cells as the anchor.

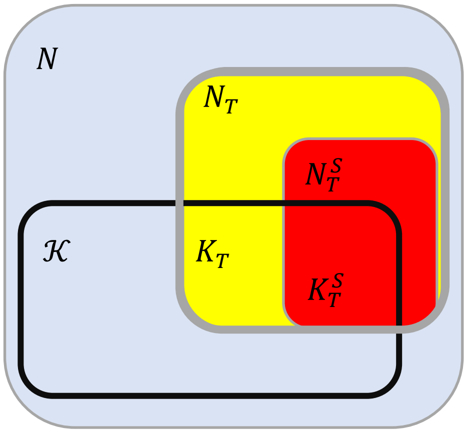


**Fig. S25.**

The enrichment score (ES) quantifies the association or exclusion of selected cells of specific cell type in the spatial proximity of the anchor cell. The variables $N$ is the number of all cells in the image, $K$ is the number of all cells in the nearest neighborhood of the anchor cell, $N_T$ is the number of all cells of cell type T in the image, $K_T$ is the number of cell type T in the nearest neighborhood of the anchor cell, $N_T^S$ is the number of all selected cells of type T and $K_T^S$ is the number of selected cell type T in the nearest neighborhood of the anchor cell.
